# Fine-tuning of the Smc flux facilitates chromosome organization in *B. subtilis*

**DOI:** 10.1101/2020.12.04.411900

**Authors:** Anna Anchimiuk, Virginia S. Lioy, Anita Minnen, Frédéric Boccard, Stephan Gruber

## Abstract

SMC complexes are widely conserved ATP-powered loop extrusion motors indispensable for the faithful segregation of chromosomes during cell division. How SMC complexes translocate along DNA for loop extrusion and what happens when two complexes meet on the same DNA molecule is largely unknown. Revealing the origins and the consequences of SMC encounters is crucial for understanding the folding process not only of bacterial, but also of eukaryotic chromosomes. Here, we uncover several factors that influence bacterial chromosome organization by modulating the probability of such clashes. These factors include the number, the strength and the distribution of Smc loading sites, the residence time on the chromosome, the translocation rate, and the cellular abundance of Smc complexes. By studying various mutants, we show that these parameters are fine-tuned to reduce the frequency of encounters between Smc complexes, presumably as a risk mitigation strategy. Mild perturbations hamper chromosome organization by causing Smc collisions, implying that the cellular capacity to resolve them is rather limited. Altogether, we identify mechanisms that help to avoid Smc collisions and their resolution by Smc traversal or other potentially risky molecular transactions.

## Introduction

Members of the family of SMC proteins are ubiquitous in eukaryotes and also present in most bacteria and some lineages of archaea. They are crucial for establishing 3D genome organization inside cells, laying the foundation for faithful segregation, recombination and repair of the chromosomal DNA molecules. Together with kleisin and kite subunits (or kleisin and hawk subunits), SMC proteins form ATP-hydrolyzing DNA motors which actively fold chromosomal DNA molecules apparently by DNA loop extrusion (Yatskevich et al., 2019). Loop extrusion can explain diverse folding phenomena across all domains of life: formation of topologically associated domains (TADs) in interphase, lengthwise compacted chromosomes during mitosis, as well as juxtaposition of the arms of bacterial chromosomes.

Recently, ATP-dependent loop extrusion has been recorded in single molecule experiments. Purified yeast condensin and vertebrate cohesin extrude DNA loops at rates of ~1 kb/s in an asymmetric (one-sided) or symmetric (two-sided) manner, respectively (Davidson et al., 2019; Ganji et al., 2018; Y. Kim et al., 2019). Nevertheless, the molecular underpinnings of loop extrusion are yet to be discovered. In the case of yeast condensin, two DNA-loop-extruding complexes were reported to occasionally traverse one another *in vitro* thus forming interlinking loops (also termed Z loops) (E. Kim et al., 2020). In principle, such behavior could improve the otherwise poor loop coverage achieved by one-sided motors but on the other hand it likely generates undesirable DNA entanglements (such as pseudoknots) (Banigan et al., 2020). The relevance of Z-loop formation and condensin traversal is yet to be determined.

Two distinct patterns of chromosome organization have been described for bacteria. One appears to rely on MukBEF (or MksBEF) complexes, presumably starting DNA loop extrusion from randomly chosen entry sites on the bacterial chromosome (Lioy et al., 2018, 2020; Mäkelä & Sherratt, 2020). In most bacteria, Smc-ScpAB complexes start DNA translocation from predefined entry sites, the 16-bp *parS* DNA sequences, which are generally found near the replication origin and are specifically recognized by the Smc-loader protein ParB (Gruber & Errington, 2009; Sullivan et al., 2009). ParB dimers form DNA clamps that self-load onto DNA at a *parS* site (Jalal et al., 2020; Osorio-Valeriano et al., 2019; Soh et al., 2019). As Smc complexes translocate away from the *parS* loading site in both directions (two-sided), they co-align the left and the right chromosome arms that flank the replication origin (Minnen et al., 2016; Tran et al., 2017; Wang et al., 2015, 2017). Bacterial genomes often have one or few closely positioned *parS* sites (separated by a few kb) (Livny et al., 2007). *B*. *subtilis* (*Bsu*), however, harbors eight *parS* sites scattered over a much wider region of the genome (~0.75 Mb) (Figure 1A).

**Figure 1.**
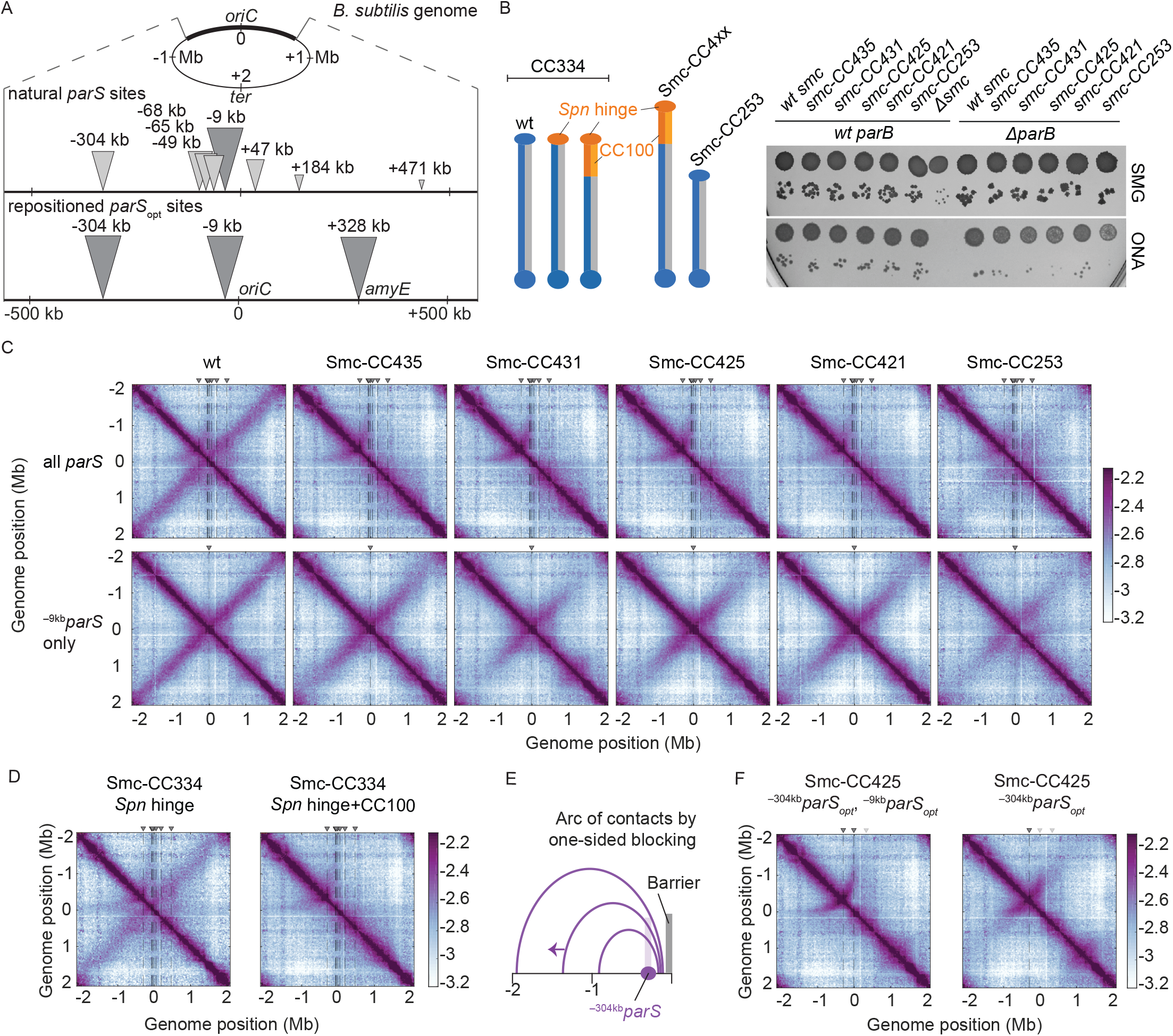
Arm-modified Smc proteins fail to align chromosome arms unless most *parS* sites are removed. A. *Upper panel*, scheme depicting the distribution of *parS* sites on the *Bacillus subtilis* genome. Triangles indicate positions of *parS* sites, size of which was scaled according to ParB occupancy as judged by ChIP-seq results (Minnen et al., 2016). *Lower panel*, scheme depicting *parS* modifications in subsequent experiments. *parS* sites were either eliminated by mutation or substituted for the *parS*_opt_ sequence (i.e. the sequence of ^−9kb^*parS*) as needed. For some experiments, an additional site (^+328kb^*parS*_opt_) was created by insertion of the *parS*_opt_ sequence into the *amyE* locus. B. *Left panel*, scheme of Smc coiled coil variants investigated in the study: wild-type (CC334), elongated (CC4xx) and shortened (CC253). *Spn* hinge+CC100, *Streptococcus pneumoniae* hinge domain and 100 amino acids hinge-proximal coiled coil (in orange colours). The coiled coil was shortened or elongated starting from a chimeric protein having the *Bsu* Smc hinge domain replaced by the *Streptococcus pneumoniae* (*Spn*) Smc hinge domain including a ~100 aa (amino acid) segment of the adjacent coiled coil. *Right panel*, spotting assay of strains with resized Smc coiled coil in wild-type or sensitised background (Δ*parB*). 9^2^ and 9^5^ dilutions were spotted on nutrient-poor (SMG) or nutrient-rich medium (ONA) and imaged after 36 hr and 15 hr, respectively. Note that in the absence of ParB, the ParABS system is non-functional and Smc loading is inefficient and untargeted, together putting a strain on chromosome segregation (Minnen et al., 2016; Wilhelm et al., 2015). The expression levels of some of these constructs (CC435, CC253) were previously shown to be close to the levels of the wild-type protein by immunoblotting (Bürmann et al., 2017). C. Normalized 3C-seq contact maps obtained from exponentially growing cultures. *Top row*, strains with wild-type *parS* sites. *Bottom row*, strains with a single ^−9kb^*parS*_opt_ (*par-S359*) site. All 3C-seq maps presented in this study are split into 10 kb bins and have the origin placed in the middle. The interaction score is in log_10_ scale, the darker the colour, the more interactions between given loci (see Materials and Methods). D. Normalized 3C-seq contact maps obtained from exponentially growing cultures carrying all the wild type *parS* sites and restored wild type length Smc (Smc-CC344) with either only hinge replaced (*Spn* hinge, *left panel*) or hinge with 100 amino acids of hinge-proximal coiled coil (*Spn* hinge + CC100, *right panel*). E. Scheme for asymmetric loop extrusion starting at ^−304kb^*parS* (*parS-334*) due to blockage of translocation towards the replication origin by head-on encounters (with other Smc complexes) generating an arc of contacts in the 3C-seq maps. F. Normalized 3C-seq contact maps of elongated Smc (Smc-CC425) carrying *parS*_opt_ sites at −304 kb and −9 kb (*left panel*) or *parS*_opt_ site at −304 kb only. Triangles above the contact map point to positions of *parS* sites (dark triangles indicate active parS sites, light triangles for reference are *parS* sites not used in given experiment).

The two-sided DNA translocation by Smc-ScpAB has two main functions: (i) it organizes bacterial chromosomes by co-aligning chromosome arms as mentioned above and (ii) it supports chromosome individualization presumably by localizing knots and precatenanes (*i*.*e*. DNA intertwinings) on the replicating chromosome, thus enabling DNA topoisomerases to completely untangle nascent sister DNA molecules efficiently (Bürmann & Gruber, 2015; Orlandini et al., 2019; Racko et al., 2018). This activity might be shared with condensin in eukaryotes (Dyson et al., 2020). The degree of defects in chromosome segregation caused by *smc* deletion is variable among species. In *B. subtilis*, chromosome segregation fails completely in *smc* mutants under nutrient-rich growth conditions but not when cells are grown with limited nutrient availability (Gruber et al., 2014; Orlandini et al., 2019; Wang et al., 2014). Deletion of *parB* or removal of *parS* sites eliminate chromosome arm alignment, but have only a mild impact on chromosome segregation, demonstrating that chromosome arm alignment is not required for efficient chromosome segregation (and for cell viability) (Lee et al., 2003; Wang et al., 2015) and implying that Smc-ScpAB can use non-*parS* sequences for loading in the absence of ParB/*parS*. DNA loop extrusion by SMC complexes may have initially arisen to facilitate individualization of sister chromosomes. Chromosome arm alignment might be an indirect—albeit putatively beneficial—consequence of two-sided loop extrusion starting from defined entry sites.

SMC complexes share a characteristic elongated architecture: a globular head and a hinge domain are connected by a long intramolecular antiparallel coiled coil ‘arm’ (Figure 1B). The functioning of the complex is restricted to discreet lengths of the coiled coil, the same periodicity of which is observed across diverse species (Bürmann et al., 2017). Two SMC proteins dimerize at the hinge and are bridged at the head domains by a kleisin subunit. This generates annular tripartite SMC-kleisin assemblies that entrap chromosomal DNA double helices (Gligoris et al., 2014; Wilhelm et al., 2015). The kite subunit (ScpB in *Bsu*) also forms dimers that associate with the central region of kleisin (ScpA in *Bsu*) (Bürmann et al., 2013). To support the nearly complete alignment of chromosome arms, Smc complexes must keep translocating on the very same DNA molecule (*i*.*e*. remain in *cis*) for the entire cycle of Smc loading, translocation and unloading (up to about 50 minutes in *B. subtilis* at 37°C). The processivity might rely on the stable entrapment of DNA double helices by the SMC-kleisin ring guarantying in-cis residence time on a given DNA molecule (Gligoris et al., 2014; Wilhelm et al., 2015). The extended nature of the coiled coils would nonetheless permit the SMC-kleisin ring to overcome relatively big obstacles (~30 nm) without necessarily dissociating from DNA. How stable DNA entrapment might be compatible with the bypass of even larger obstacles on the chromosome remains an open question (Brandão, Wang, et al., 2019). Moreover, Smc complexes loaded simultaneously at different *parS* sites (which are widely scattered in *B. subtilis*, Figure 1A) will translocate towards each other and eventually collide. Smc traversal (as proposed for the formation of Z-loops by condensin *in vitro*) could resolve such putative encounters. On the other hand, the traversal by Smc dissociating from and re-associating with DNA may result in Smc binding *in trans* thus creating persistent physical linkages between sister chromosomes rather than helping to resolve them.

Here, we studied the effect of Smc, ParB and *parS* alterations on chromosome organization to explore how Smc-ScpAB load and translocate on a chromosome with multiple loading sites. Based on our results, we propose that Smc complexes rarely meet on the chromosome under physiological conditions. We argue that multiple parameters are fine-tuned in such a way to avoid Smc-Smc collisions in the first place. Few Smc complexes are available for loading, because most Smc complexes are associated with the chromosome arms for an extended period of time. Artificially increasing the rate of encounters by mildly elevating the cellular levels of Smc complexes *in vivo*, or by reducing the chromosome residence time or translocation rate, lead to obvious perturbations in chromosome architecture, presumably due to unresolved Smc-Smc collisions. Also, the genomic clustering of strong *parS* sites seems to play a vital role in avoiding Smc collisions in *B. subtilis*. Although we cannot exclude the possibility of dedicated mechanism for the resolution of collisions *per se*, we suggest that an optimized Smc flux helps to eschew such events, presumably to avoid complications emerging from any attempted resolution reaction.

## Results

### Arm-modified Smc proteins fail to juxtapose chromosome arms

We previously isolated chimeric Smc proteins with elongated and shortened coiled coils that can functionally substitute for the *B. subtilis* Smc (Bürmann et al., 2017). From a collection of 20 constructs, we here identified several elongated Smc proteins, including Smc-CC425 (with a 425 aa coiled coil as compared to the 334 aa in wild-type Smc), which supported normal growth on nutrient-rich medium even in a sensitized background (Δ*parB*) (Figure 1B, S1D).

We hypothesized that the coiled coil length may influence DNA translocation, particularly when Smc complexes meet and overcome obstacles on the DNA track. To address this, we performed 3C-seq analysis on cells grown in nutrient-poor medium (SMG) at 37°C, supporting growth with a generation time of ~ 60 minutes. Encounters between translocating Smc complexes and the replication fork are expected to be rare under these conditions as replication initiates only about every 60 minutes (Gruber et al., 2014). We found that Smc-CC425 and the other elongated variants failed to support normal chromosome organization (Figure 1C). As revealed by the absence of a secondary diagonal, the co-alignment of chromosome arms was strongly compromised. An arc of contacts on the left arm of the chromosome however was observed in wild-type and mutant 3C-seq maps (see below). A control chimeric protein with wild-type arm length (*Spn* hinge + CC100) showed similar growth behavior and 3C-seq maps as the resized variants (Figure 1D, S1A), implying that the defect in chromosome organization was indirectly caused by the Smc arm modifications. With a shortened Smc protein, Smc-CC253, only residual levels of inter-arm contacts were noticeable, extending only few hundred kb from the replication origin (Figure 1C). We conclude that in contrast to wild-type Smc, engineered Smc variants are unable to properly co-align the two chromosome arms despite supporting growth, and presumably chromosome segregation also, apparently normally.

### Removal of all but one *parS* sites rescues chromosome folding by arm-modified Smc

To reveal the cause of the defect in chromosome arm alignment, we sought to characterize the loading and translocation of the modified Smc complexes on the bacterial chromosome. We started by generating strains in which seven *parS* sites were inactivated by mutations, with ^−9kb^*parS* (*parS-359*) remaining the only *parS* site on the chromosome (along with the weak ^+1058kb^*parS* site; *parS-90*; see below). As expected from published work, wild-type Smc efficiently aligned chromosome arms from a single strong *parS* site (Wang et al., 2017) (Figure 1C). Curiously, all four Smc proteins with an extended Smc arm displayed clearly increased levels of inter-arm contacts (Figure 1C). Near the replication origin, chromosome arm alignment was comparable to wild type, while the inter-arm contacts were less frequent (or absent) further away from the replication origin with all modified Smc constructs (Figure 1C). The shortened Smc variant (CC253) also displayed more inter-arm contacts when the seven *parS* sites were mutated. Thus, the removal of *parS* sites improved—rather than hindered— chromosome arm alignment by modified Smc proteins.

The arc of contacts detected on the left arm of the chromosome was lost in all strains harboring only the ^−9kb^*parS* site (Figure 1C) (Marbouty et al., 2015). It was also lost when only ^−304kb^*parS* site (*parS-334*) was mutated (Figure S1C). Of note, the ^−304kb^*parS* site is unique, in being relatively strong as well as distantly located from other strong *parS* sites (Figure 1A). DNA loop extrusion starting from this site is asymmetric, presumably due to the high likelihood of a clash with other Smc complexes (Figure 1E) (see below).

To test the impact of *parS* distribution in a more controlled way, we created strains with two distantly positioned *parS* sites (Figure 1A, lower panel). Since *parS* sites accumulate varying levels of ParB protein (Figure 1A) (Graham et al., 2014; Minnen et al., 2016), we first identified the *parS* sequences giving highest chromosomal recruitment of ParB and Smc when inserted at the *amyE* locus (+ 328 kb) in otherwise wild-type cells. The sequence of the ^−9kb^*parS* site outperformed four other natural *parS* sequences as well as an engineered consensus sequence at the ectopic location as judged by ChIP-qPCR (Figure S2A). We thus used the strong ^−9kb^*parS* sequence (denoted as *parS*_opt_) in subsequent experiments. When two *parS*_opt_ sites (^−9kb^*parS*_opt_ and ^−304kb^*parS*_opt_) were combined on the same chromosome, chromosome arm alignment by Smc-CC425 became inefficient, producing a contact map similar to the one obtained with all *parS* sites present (Figure 1F). We conclude that the presence of two or more *parS* sites hampers chromosome organization by Smc-CC425, conceivably because Smc-CC425 is more prone to collisions than wild-type Smc or less efficient in resolving them.

### An arm-modified Smc protein over-accumulates in the replication origin region

Wild-type Smc-ScpAB displays highest enrichment on the chromosome in the replication origin region with long and shallow gradients of enrichment along both chromosome arms (Gruber & Errington, 2009; Minnen et al., 2016), presumably generated by loading at *parS*, by translocation towards the replication terminus (*ter*), and by rare and stochastic unloading from the chromosome arms. Removal of seven *parS* sites had only a minor impact on the distribution of wild-type Smc-ScpAB as judged by chromatin immunoprecipitation coupled to deep sequencing (ChIP-seq) using α-ScpB serum (Figure 2A, B, left panels). The chromosomal distribution of Smc-CC425 was markedly different (Figure 2A). It showed hyper-enrichment near the replication origin and poor distribution towards the chromosome arms, indicating that the modified Smc coiled coil impeded DNA translocation and/or increased the rate of unloading. Remarkably, the removal of seven *parS* sites substantially reduced the hyper-enrichment near the origin and increased the otherwise low signal on the chromosome arms (Figure 2A, B, right panels, S2B). The hyper-enrichment of Smc in the replication origin region thus correlated with the loss of chromosome arm alignment (Figure 1C) in these strains. We next synchronized chromosomal loading of Smc and Smc-CC425 at a single *parS* site (^−9kb^*parS*_opt_) in a population of cells by depleting and repleting ParB protein. These experiments were performed at 30°C to allow sufficient ParB expression from a theophylline riboswitch-regulated *parB* construct. Smc and Smc-CC425 complexes were found enriched in a ~700 kb region centered on the replication origin after 20 minutes of ParB induction by ChIP-seq analysis using α-ScpB serum (Figure 2C, S2C). For wild-type Smc, the enriched region increased in size over time, inferring a constant DNA translocation rate of roughly 500 bp/sec at 30°C. Notably, the high enrichment near *parS* disappeared at the later time points as Smc-ScpAB became broadly distributed on the chromosome. The region of Smc-ScpAB enrichment also broadened in Smc-CC425 during the second and third time interval, albeit with an apparently reducing rate. In addition, the origin region remained highly enriched in ScpB also at the later time points. These two observations suggest that the Smc-CC425 protein has a reduced chromosome residence time and/or a reduced translocation rate. Determining a meaningful translocation rate for Smc-CC425 is difficult because of a wide gradient distribution at the later time points. Using 3C-seq, we observed that Smc-CC425 was able to align chromosome arms in this experimental system, yet the alignment did not extend all the way to the terminus region (Figure 2D). Moreover, the onset of chromosome alignment as well as the rate of progress were somewhat reduced when compared to wild-type Smc. These experiments demonstrate that Smc-CC425 efficiently accumulated in the replication origin region, but the redistribution to the chromosome arms, particularly to distal loci, was hampered.

**Figure 2.**
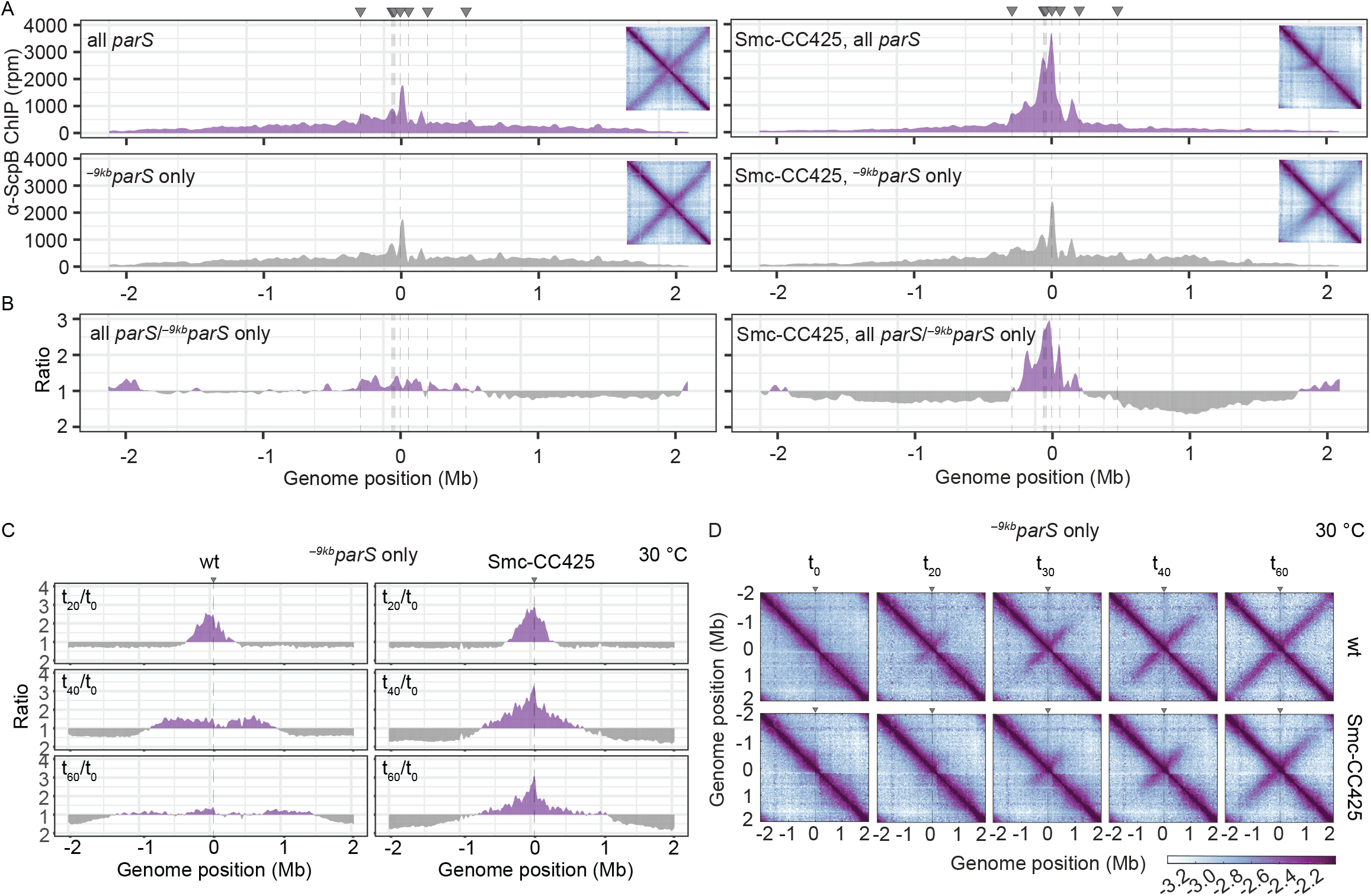
Modified Smc proteins hyper-accumulate in the replication origin region. A. Read count distribution for chromatin immunoprecipitation coupled to deep sequencing (ChIP-seq) using α-ScpB serum. *Left panel*, strain carrying wild-type Smc with wild-type *parS* sites (*top*) or single ^−9*kb*^parS_opt_ (*parS-359*) site (*bottom*). Removal of *parS* sites results in a slightly reduced enrichment in the origin region and in turn modestly increased signal mainly on the right arm of the chromosome (supposedly due to presence of the weak ^+1058kb^*parS* site; *parS-90*). *Right panel*, strain carrying Smc with elongated coiled coil (Smc-CC425) with wild-type *parS* sites (*top*) or single ^−9kb^parS_opt_ (*parS-359*) site (*bottom*). Insets depict corresponding 3C-seq contact maps. All ChIP-seq profiles presented in this study are divided into 1 kb bins and have the origin placed in the middle. Dashed lines indicate the position of *parS* sites. B. Ratio plots of respective ChIP-seq experiments for wild-type and elongated Smc (Smc-CC425). For each bin, normalized read counts were compared with respective wild-type values. The higher value was divided by the lower. If the mutant/wild-type ratio was > 1, it was plotted above the genome position axis (in violet colours). If the wild-type/mutant ratio was > 1, the ratio was plotted below the axis (in grey colours). C. ChIP-seq time course experiments using α-ScpB serum for strains carrying wild-type (*left panel*) or elongated Smc (Smc-CC425, *right panel*). Strains harbour a single loading site, ^−*9kb*^*parS*_opt_ (*parS-359*), and theophylline-inducible *parB* gene. Ratios of given time point (t_x_) versus t_0_ are shown. For each bin, normalized read counts were compared with respective t_0_ value and the higher value was divided by the lower. If the ratio t_x_/t_0_ was > 1, it was plotted above the genome position axis (in violet colours). If the ratio t_0_/t_x_ was > 1, the ratio was plotted below the axis (in grey colours). D. Normalized 3C-seq contact maps for the time course experiments with strains carrying wild-type (*top panel*) or elongated Smc (Smc-CC425, *bottom panel*) as in (C).

A simple explanation for the hyper-accumulation of Smc-CC425 in the replication origin region is an increased unloading rate, possibly in combination with a reduced translocation rate particularly in the presence of multiple *parS* sites. With less time spent translocating along the chromosome arms, the cytoplasmic pool of Smc increases and as a consequence so does the flux of loading, which, together with a reduced translocation rate, leads to artificially increased enrichment near the *parS* site(s). This in turn elevates the probability of Smc-Smc collisions on the chromosome when loading occurs at multiple *parS* sites but not when restricted to a single *parS* site. Collisions further exacerbate the Smc hyper-enrichment by hindering Smc translocation away from *parS* sites (Figure 2A, right panel). Reduced chromosome residence times and/or reduced translocation rates thus explain all phenotypic consequences of the Smc arm-modifications. Whether Smc-CC425 also has a problem in resolving collisions remains to be established (see discussion).

### Wild-type Smc protein generates overlapping chromosome folding patterns

We next wondered how wild-type Smc proteins co-align chromosome arms when starting DNA loop extrusion at multiple *parS* sites. Wild-type Smc displayed relatively low enrichment in the replication origin region even when all natural *parS* sites were present (Figure 2A). To understand how collisions between translocating Smc complexes are avoided or resolved, we next aimed to increase the incidence of collisions by positioning two *parS*_opt_ sequences at selected sites in varying genomic distances and performed 3C-seq analysis.

As expected, control strains with a unique *parS*_opt_ sequence at positions −9 kb, at −304 kb, or at +328 kb (at *amyE*) demonstrated extensive alignment of the respective flanking regions (Figure 3A) (Wang et al., 2015, 2017). The ^+328kb^*parS*_opt_ site resulted in asymmetric alignment of the flanking DNA probably due to the presence of clusters of highly transcribed genes (including rDNA operons) in head-on orientation (Brandão, Paul, et al., 2019). Of note, the frequency of contacts reaching beyond the replication origin is notably reduced with ^−^304kb*parS*_opt_ or ^+328kb^*parS*_opt_, implying that the origin region acts as a semi-permissive barrier for Smc translocation, as previously indicated (Minnen et al., 2016), (or a Smc unloading site).

**Figure 3.**
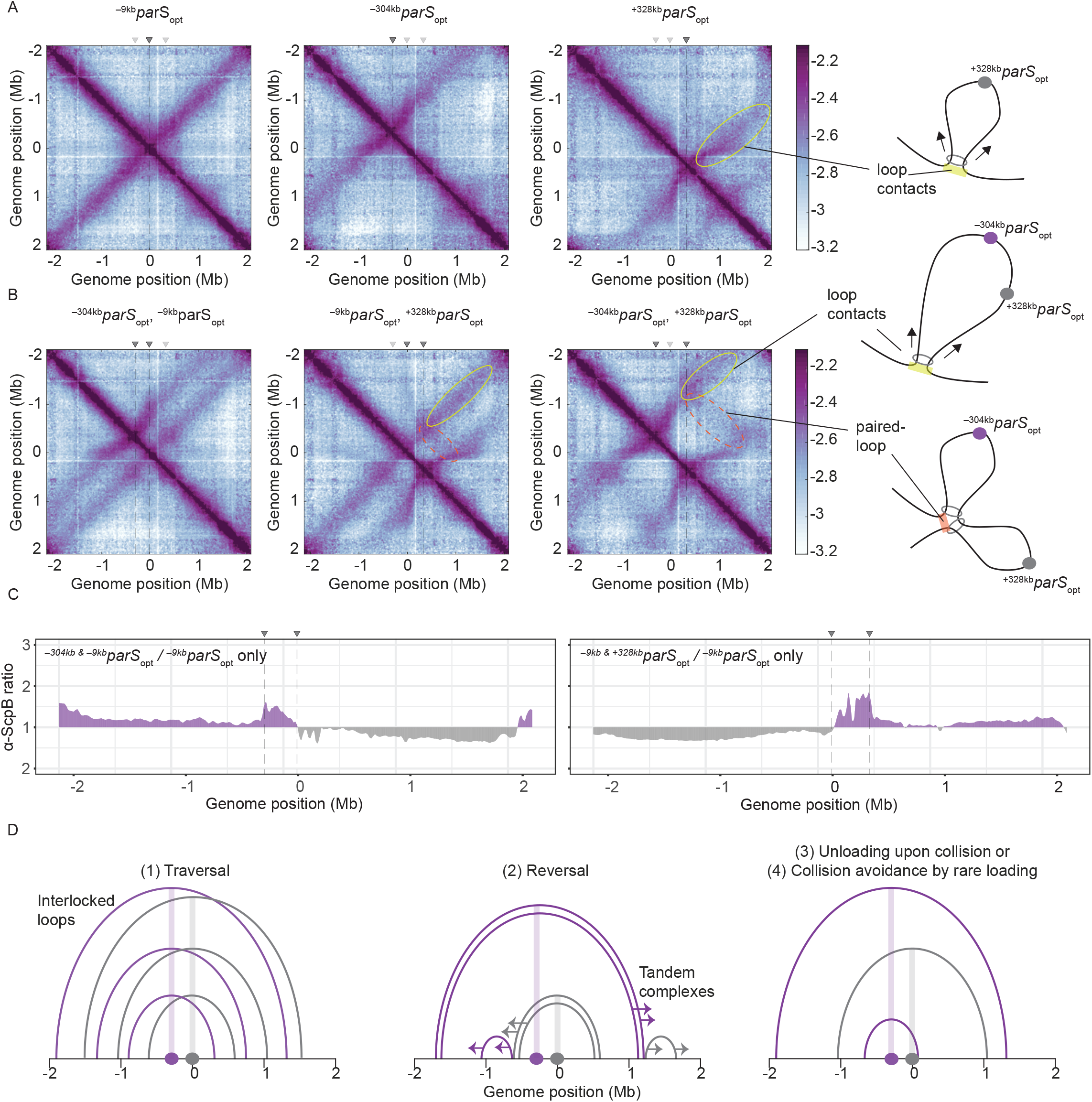
Overlapping chromosome arm alignment patterns for wild type Smc. A. Normalized 3C-seq contact maps for strains with a single *parS*_*opt*_ site at −9 kb, −304 kb, or +328 kb. Schemes depict exemplary chromosome arm alignment inflicted upon a single loading site (yellow), here ^+328*kb*^parS_opt_. Dark grey triangles above the contact maps indicate the active *parS* site positions. B. Normalized 3C-seq contact maps for strains with two *parS*_*opt*_ sites spaced by ~300 kb (*left* and *middle*) or ~600 kb (*right*). Schemes interpreting interactions in the contact maps: loop contacts (in yellow colours) and paired-loop contacts presumably caused by colliding SMC loaded at both *parS* sites (in orange colours). C. Ratio plots for ChIP-seq read counts for a strain with two *parS* sites (left panel: ^−304*kb*^parS_opt_ and ^−9*kb*^parS_opt_; right panel: ^−9*kb*^parS_opt_ and ^+328*kb*^parS_opt_) and a control strain with a single *parS* site (^−9*kb*^*parS*_opt_). Representation as in Figure 2B. D. Schemes depicting possible scenarios for collision avoidance and collision resolution: SMCs traversal (*1*), reversal (*2*) or avoid interaction by unloading upon collision or by rare loading (*3*)(*4*).

More importantly, when two *parS* sites were combined on the chromosome, striking novel patterns of chromosome organization by wild-type Smc arose (Figure 3B). In all cases, parallel secondary diagonals emerging from the two *parS* sites were detected. The pattern observed with ^−*304kb*^*parS*_opt_ and ^−*9kb*^*parS*_opt_ can—to a large degree—be explained as a combination of contacts observed in strains with the corresponding single *parS* sites, however, with clearly reduced probability for contacts extending beyond the region demarcated by the *parS* sites. The latter appears to suggest that *parS* sequences might act as loading and unloading sites. A small but noticeable fraction of Smc complexes however managed to translocate towards and beyond other *parS* sites apparently mostly unhindered. The contact maps involving the ^−304kb^parS_opt_ and ^+328kb^*parS*_opt_ sites showed additional contacts likely representing paired loops originating from collided Smc complexes loaded at opposite *parS* sites (Figure 3B). The presence of such paired loop contacts was less clear for the other *parS* combinations possibly due to background signal and limited resolution of the 3C-seq maps. We conclude that wild-type Smc-ScpAB complexes rarely block one another when loaded from all natural *parS* sites (with the notable exception of ^−304kb^*parS*). When the distance between two strong *parS* sites was artificially increased, however, impacts arising from collisions and blockage became noticeable. The blockage of Smc translocation was also apparent from ChIP-seq analysis, which demonstrated over-enrichment of Smc between two *parS* sites (^−304kb^*parS*_opt_ and ^−9kb^*parS*_opt_ or ^−9kb^*parS*_opt_ and ^+328kb^*parS*_opt_) when compared to the single *parS* control (Figure 3C, S3B, S3C). The effects of collisions on chromosome organization and Smc distribution are thus subdued but detectable with wild-type Smc.

To explain the relatively mild impact of collisions in wild-type cells, we envisaged following scenarios: (1) the traversal of Smc complexes generating interlocking loops (Brandão et al., 2020; E. Kim et al., 2020) (2) the reversal of the translocation of one Smc complex by opposing complexes (E. Kim et al., 2020), (3) the unloading of one or both complexes upon collision, or (4) collision avoidance either by infrequent loading (Figure 3D) or (5) by mutually exclusive *parS* usage (Figure S3A). The latter hypothesis seemed highly unlikely as all but one *parS* sites would have to remain inactive for extended periods of time. While all other scenarios seemed plausible and may contribute to the process of chromosome organization, without making additional assumptions only one scenario, the avoidance of encounters by infrequent loading, also provided an explanation for the strong defects in chromosome organization observed for Smc-CC425.

### Increasing the pool of Smc hampers chromosome organization

If Smc-CC425 indeed fails to juxtapose chromosome arms due to over-enrichment in the replication origin region, collisions may be rare in wild-type cells because of a high chromosome residence time and a limited pool of soluble Smc complexes, resulting in a small flux of Smc onto the chromosome. If so, artificially increasing the flux of Smc should lead to defects in chromosome organization with multiple *parS* sites but not with a single *parS* site (assuming that most Smc is loaded at *parS* sites) (as observed for Smc-CC425 under normal expression levels). If Smc complexes, however, were to efficiently traverse, reverse or unload one another, then increased Smc levels would not result in defective translocation and chromosome organization.

To test this prediction, we first slightly increased the cellular level of all subunits of the Smc complex by inserting an additional copy of the *smc* gene and of the *scpAB* operon under the control of their respective endogenous promoters into the genome. The increased levels of Smc-ScpAB did not noticeably affect cell growth (Figure S4A). Immunoblotting suggested a 4-5-fold increase in Smc and ScpB protein levels in the SMC^high^ strain when compared to wild type (Figure 4A). Next, we performed 3C-seq analysis. Chromosome arm co-alignment was strongly hampered—rather than improved—by the presence of extra Smc complexes in the cell (Figure 4B). A prominent arc was formed at the position of the ^−304kb^*parS* site and the secondary diagonal originating in the origin region was weak and diffuse in the SMC^high^ strain. This defect was fully restored, however, by removal of seven *parS* sites (with the remaining strong site being either ^−9kb^*parS*_opt_ or ^−304kb^*parS*_opt_) (Figure 4C). Note that an additional feature (a minor secondary diagonal) present on the right arm of the chromosome likely originated from Smc loading at the weak ^+1058kb^*parS* site. The presence of two strong *parS* sites (^−9kb^*parS*_opt_ and ^−304kb^*parS*_opt_) led to a new pattern of chromosome folding in the SMC^high^ strain. The alignment of DNA flanking the *parS* sites became highly asymmetric (Figure 4D).

**Figure 4.**
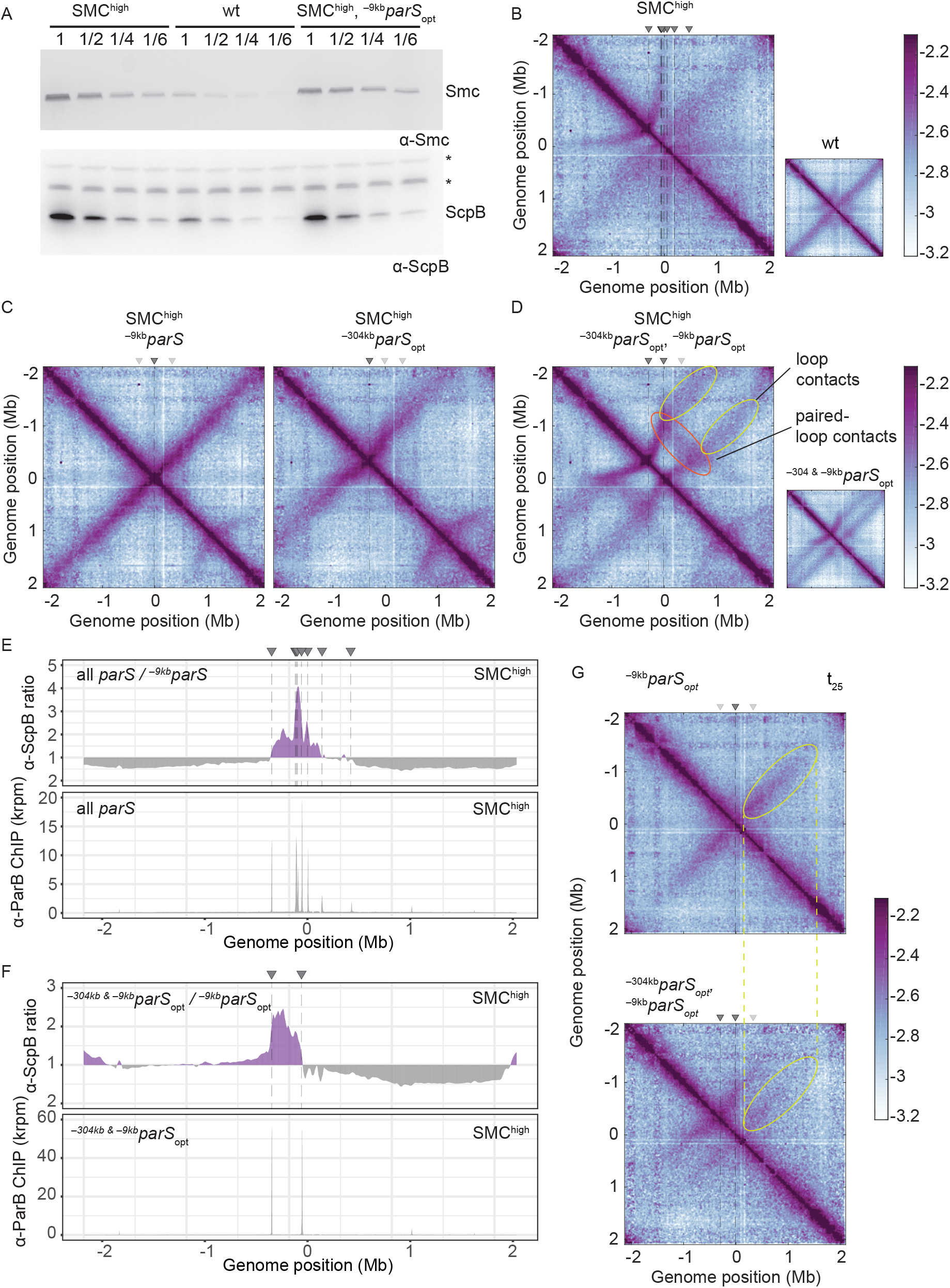
Increasing the cellular pool of Smc hampers chromosome organization. A. We estimated the relative increase in Smc and ScpB abundance in cells of this ‘SMC^high^’ strain by immunoblotting using α-Smc and α-ScpB serum, respectively. SMC^high^ denotes strains with extra genes. Protein extracts of wild-type or SMC^high^ strains (harboring all *parS* sites or single *parS* site) were serially diluted with extracts from *Δsmc* or *ΔscpB* strains as indicated (see Materials and Methods). * indicates unspecific bands generated by the α-ScpB serum. B. Normalized 3C-seq contact map for SMC^high^ strain with all *parS* sites present. Inset shows 3C-seq contact map of a strain with wild-type protein levels (also displayed in Figure 1C) for direct comparison. C. Normalized 3C-seq contact maps for SMC^high^ strains with *parS*_*opt*_ at −9 kb only or at −304 kb only. D. Normalized 3C-seq contact map for SMC^high^ strain with *parS*_*opt*_ at positions: −9 kb and −304 kb. As in (B), with inset displaying respective control strain with normal Smc expression (also shown in Figure 3B). E. Ratio plots for ChIP-seq read counts comparing SMC^high^ strains with all *parS* sites and a single *parS* site (^−9*kb*^parS_opt_). Representation as in Figure 2B (*top panel*). Read count for α-ParB ChIP-seq in SMC^high^ strain (*bottom panel*). F. As in (E) involving a SMC^high^ strain with two *parS* sites (^−304*kb*^*parS*_opt_ and ^−9*kb*^*parS*_opt_) instead of all *parS* sites. G. Normalized 3C-seq contact maps for time point t_25_ after IPTG-induced ParB expression with a single *parS*_*opt*_ site (*top*) or two *parS*_*opt*_ sites (at −9kb and −304kb) (*bottom*). Ovals (in yellow colours) mark the position of contacts stemming from loop extrusion originating at ^−^9k*bparS*_*opt*_.

Moreover, the contacts corresponding to paired loops became clearly visible (Figure 4D). Finally, contacts outside the *parS*-demarcated region were rare and spread out, and their center was shifted away from the *parS* sites. The latter indicated that Smc complexes that were loaded at one *parS* site and managed to move beyond the other *parS* site have experienced a strongly reduced translocation rate from one to the other *parS* site, presumably due to encounters with and temporary (or partial) blockage by Smc complexes translocation in opposite orientation.

If so, then extra Smc levels may lead to artificially higher accumulation of Smc-ScpAB near the replication origin when multiple *parS* sites are present but not with a single *parS* site, as observed for the modified Smc at normal levels of expression (Figure 2A). To test this, we performed ChIP-seq with α-ParB and α-ScpB serum in SMC^high^ strains. The α-ParB ChIP-seq demonstrated that the localization of ParB to *parS* sites is, as expected, largely unaffected by the increased levels of Smc (Figure 4E) (Minnen et al., 2016). The chromosomal distribution of ScpB was also largely unaffected in SMC^high^ cells harboring a single ^−*9kb*^parS_opt_ site (Figure S4B). However, in the presence of two or multiple additional *parS* sites, the enrichment between the *parS* sites increased strongly (Figure 4E, F, S4C). The changes in ScpB distribution upon *parS* site removal are strikingly similar in Smc-CC425 and SMC^high^ (Figure 2B, 4E), supporting the notion that both modifications lead to more frequent collisions and blockage probably by the same mechanism: an increased rate of Smc loading.

### Synchronized Smc loading favors Smc collisions

Finally, we synchronized the loading of Smc by induction of ParB with the idea that ParB repletion transiently elevates the rates of Smc loading (from a larger cytoplasmic pool of Smc) and thus increases the rates of encounters even with normal cellular levels of Smc-ScpAB. Here, we used a different inducible promoter, the IPTG-inducible *Pspank* (Wang et al., 2017), which enabled us to grow cells at 37°C and compare the results more directly to the experiment with constitutively expressed ParB (Figure 3B). We found that the alignment of DNA starting from ^−9kb^*parS*_opt_ site was indeed hampered when a second parS site, ^−304kb^*parS*_opt_, was present (Figure 4G, S4D), even more so than with continuous ParB expression (Figure 3B).

## Discussion

Establishing how SMC complexes manage to organize and orderly compact DNA in the crowded environment of a cell is a burning question in the field. SMC complexes translocate along an unusually flexible, congested and entangled translocation track, *i*.*e*. the ‘chromatinized’ DNA double helix. The architecture of SMC complexes—one of a kind amongst the collection of molecular motors—is likely a reflection of a unique translocation mechanism. To support the folding of Mb-sized regions of the chromosome, Smc complexes need to keep translocating on the same DNA double helix from initial loading to unloading (a process lasting several tens of minutes in bacteria). Assuming a topological SMC-DNA association (Gligoris et al., 2014; Wilhelm et al., 2015), staying on the translocation track *in cis* is guaranteed as long as the SMC-kleisin ring remains closed thus preventing the release of DNA(s).

During translocation, SMC complexes must frequently overcome obstacles on the DNA. Very large obstacles (> 30 nm) could not be overcome while keeping DNA entrapped but would need to be bypassed by dissociating from the translocation track transiently. Such obstacles might include branched DNA structures, protein-mediated DNA junctions (i.e. crossings in the translocation track) as well as other SMC complexes bound at the base of large DNA loops. Traversal and bypassing are not risk-free strategies. When transiently disconnecting from DNA, the complex risks the loss of directional translocation by wrongly reconnecting with the same DNA double helix or even establishes an unwanted trans-DNA linkage by connecting with a different DNA double helix. Any straying onto the sister DNA molecule (going *in trans*) would not only defeat the purpose of DNA loop extrusion but actually actively hinder chromosome segregation. Here, we addressed the balance between avoiding and resolving Smc collisions.

### Avoiding Smc encounters

In our study, we show that impacts from collisions are barely noticeable in wild-type cells. Under physiological conditions, collisions between Smc-ScpAB complexes are kept at a tolerable level by a low cellular abundance of Smc-ScpAB, a high DNA translocation rate, an extended time of residence on the chromosome arms, and the preferential usage of the origin-proximal *parS* sites (^−68kb^*parS*, ^−63kb^*parS*, ^−49kb^*parS*, ^−9kb^*parS*). Thus, multiple strategies are employed to avoid Smc encounters, possibly rendering the resolution of Smc collisions unnecessary. A rough estimate for the cellular abundance of Smc-ScpAB complexes is intriguingly low: ~30 Smc dimers per chromosome or less (i.e. about one Smc complex per ~140 kb of chromosomal DNA or less) (Wilhelm et al., 2015). With the main *parS* sites being clustered in a ~60 kb region of the genome (Figure 1A), collisions are accordingly expected to be rare (with or without traversal). Occasional loading of Smc at one of the more distal *parS* sites however would quite often lead to collisions, thus resulting in contact maps with an arc-shaped pattern, such as observed for ^−304kb^*parS* (the strongest of the distal *parS* sites) (Figure 1C), (Brandão et al., 2020). When strong *parS* sites are artificially moved further away from each other, then impacts from collisions also become obvious (Figure 3). The obvious defects in chromosome organization observed with altered *parS* positioning, elevated Smc levels or engineered Smc proteins however do not substantially impact bacterial growth, suggesting that chromosome segregation is efficiently supported even with Smc collisions (and without noticeable chromosome arm alignment). Colliding Smc complexes might thus efficiently promote DNA disentanglement (by DNA topoisomerases) but hamper the chromosome folding process, possibly leading to large cell-to-cell variations in chromosome organization with likely knock-on effects on other cellular processes including nucleoid occlusion and cell division.

The presence of multiple types of SMC complexes acting on the same chromosome likely aggravates the issue of collisions. In *Pseudomonas aeruginosa* (*Pae*), the impact of collisions seems to be dealt with by a hierarchy amongst two endogenous SMC complexes (Lioy et al., 2020). Smc-ScpAB appears to limit the loop extrusion activity of MksBEF but not *vice versa*. When the heterologous *E. coli* MukBEF complex was introduced in place of MksBEF, it blocked the activity of *Pae* Smc-ScpAB but not *vice versa*. The hierarchy is possibly given by the relative abundance of these complexes and by differing residence times on the chromosome.

### Resolving Smc encounters?

The parsimonious explanation for our observations—not requiring the involvement of dedicated and potentially hazardous molecular transactions—is that neither wild-type nor modified Smc proteins are able to traverse one another. A recent study quantitatively describing Smc action in *B. subtilis* by simulations, however, suggested that Smc traversal is needed to recapitulate the precise distribution of contacts obtained with an artificial *parS* arrangement (similar to Figure 3B) (Brandão et al., 2020). While the simulations are not fully conclusive due to substantial uncertainties concerning many involved parameters, the possible existence of unexplored alternative explanations (e.g. Figure 3D), and the lack of a molecular understanding, they seem to support the notion of interlinked DNA loops on the bacterial chromosome, which was previously proposed for yeast condensin based on experiments with purified components (E. Kim et al., 2020). A putative defect in Smc traversal might help to explain the over-accumulation of Smc-CC425 in the replication origin region and the defective chromosome organization. If so, it is tempting to speculate that the nature and the integrity of the Smc hinge domain and the adjacent coiled coil is critical for traversal. The unique donut-shaped architecture of the hinge is widely conserved, yet the hinge can be replaced by the Rad50 zinc hook without significant functional perturbations (Bürmann et al., 2017). The involvement of the hinge specifically in Smc traversal might explain its wide conservation despite it not being essential for the basic loop extrusion mechanism. Taking the notion further, a traversal-related mechanism involving the cohesin hinge domain might establish sister chromatid cohesion at the replication fork (Gruber et al., 2006; Srinivasan et al., 2018). MukBEF on the other hand might not be able to support traversal (at least using the same mechanism) as its MukB hinge is more highly diverged e.g. lacking the central channel (Ku et al., 2010). How Smc traversal might occur without risking establishing unwanted DNA linkages yet is totally unclear.

### Multiple *parS* sites on the *B. subtilis* chromosome

The presence of multiple *parS* sites likely improves the robustness of the chromosome segregation process (Böhm et al., 2020). Most bacteria have clustered *parS* sites (within 5-40 kb region) and are sensitive to deleting or dispositioning them outside a tolerance region (Böhm et al., 2020; Lagage et al., 2016; Minnen et al., 2011; Tran et al., 2017). Severe consequences of manipulating *parS* distribution include a longer generation time (when *mksBEF* is missing) (Lagage et al., 2016), an increased number of anucleate cells (Böhm et al., 2020) and elongated cells (Tran et al., 2017). Some of these defects might be related to altered Smc function. In *C. crescentus*, Smc translocates only ~600 kb away from *parS* (Tran et al., 2017). The partial chromosome arm alignment in *C. crescentus* is reminiscent of the observation with modified Smc proteins in *Bsu*. It is tempting to speculate that a shorter chromosome residence time or a lower translocation rate of Smc-ScpAB is acceptable when *parS* sites are tightly clustered or when combined with a single *parS* site. Some bacterial genomes however harbor multiple *parS* sites that are quite widely scattered on the genome. From the point of view of collision avoidance, this seems counterproductive. Intriguingly, the scattered *parS* distribution is restricted to few lineages on the phylogenetic tree of bacteria, including Bacilli (Livny et al., 2007). The scattering of *parS* sites likely serves a dedicated purpose in the lifestyles of these bacteria, in *B. subtilis* possibly during sporulation when large chunks of the genome need to be captured at the cell pole to promote entrapment of the chromosome in the pre-spore. This process might benefit from the condensation of the replication origin rather than an alignment of chromosome arms. Consistent with this notion, chromosome arm alignment is lost when DNA replication is artificially blocked (as naturally occurring during sporulation) and replaced by smaller loops formed at individual *parS* sites (Wang et al., 2015). A simple explanation for this altered pattern of chromosome folding during replication blockage (and possibly also during sporulation) would be an increased rate of Smc collisions due to an elevated (Smc) protein-to-DNA ratio.

Altogether, our results strongly suggest that the process of Smc DNA translocation is finely tuned to keep the probability of Smc encounters at a low level, presumably to enable extensive DNA loop extrusion without the need to resolve Smc collisions.

## Materials and Methods

### *Bacillus subtilis* strains and growth

*Bsu* 1A700 or PY79 isolate was used for experiments. Naturally competent *Bsu* was transformed via homologous recombination as described in (Bürmann et al., 2013) and selected on SMG-agar plates with appropriate antibiotic selection. Transformants were next checked by PCR and, if required, Sanger sequencing. Genotypes of strains used in this study are listed in Table 1.

**Table 1.**
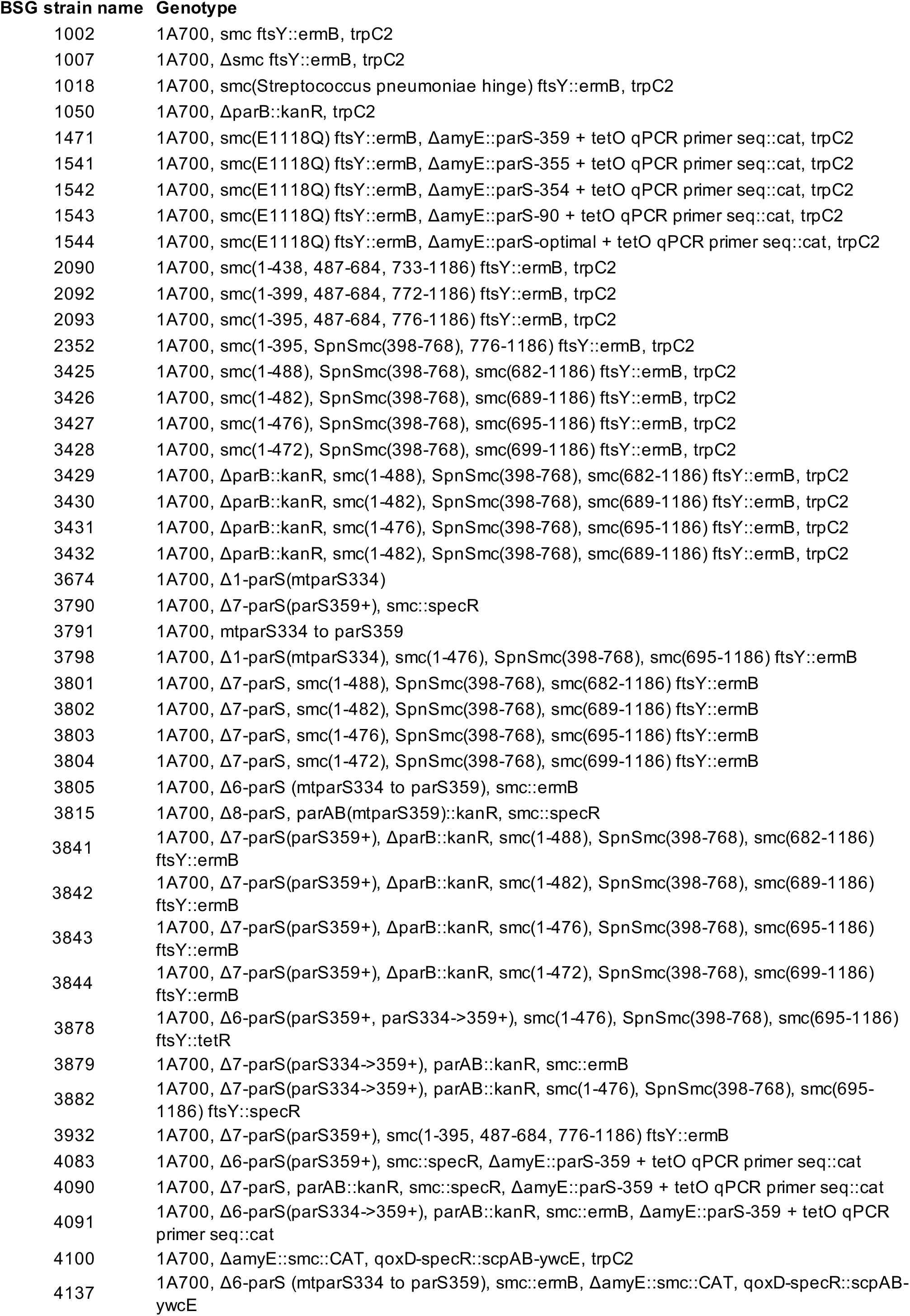
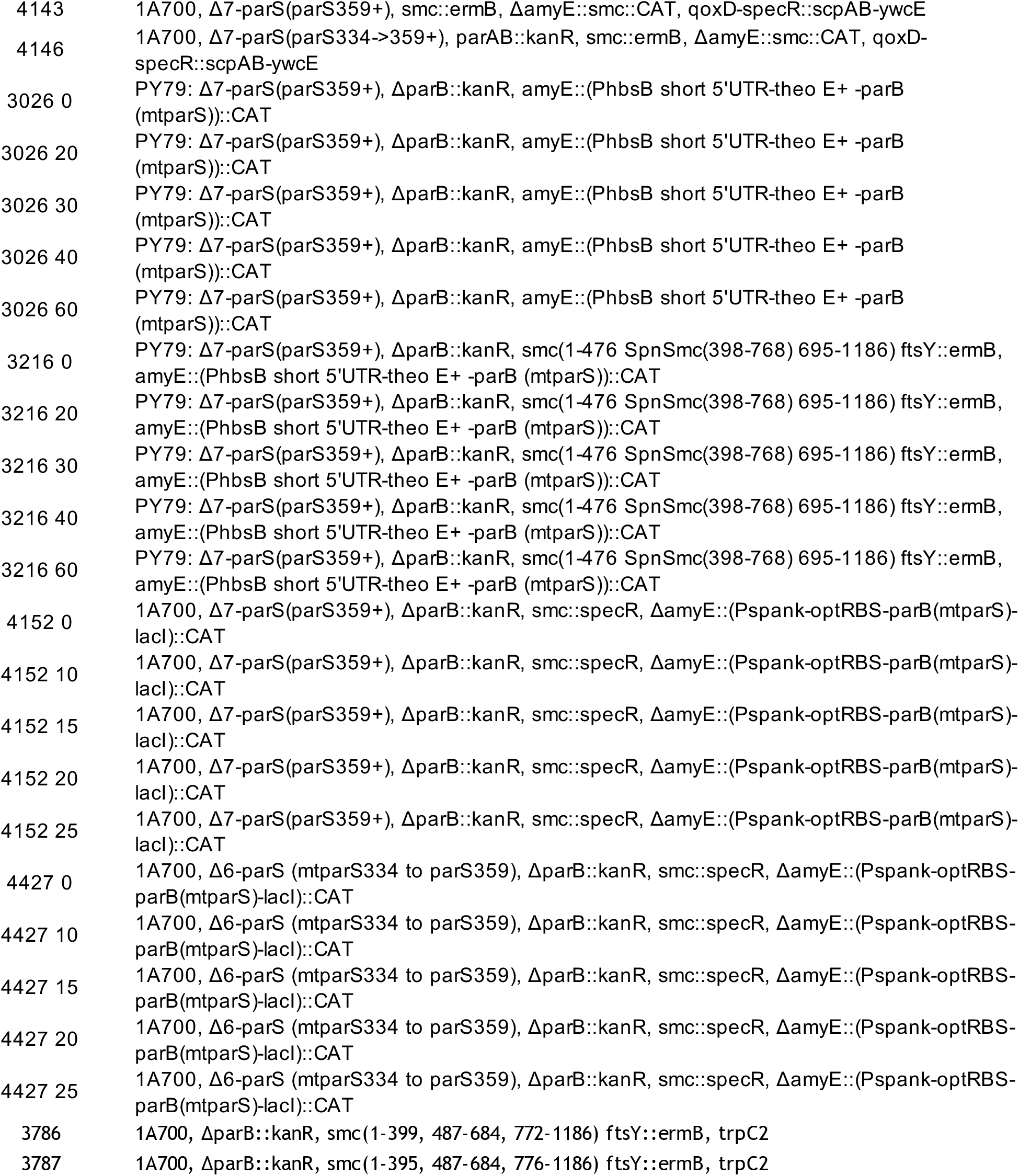
List of strains and genotypes used in the study

For spotting assays, the cells were cultured in SMG medium at 37 °C to stationary phase and 9^−2^ and 9^−5^ - fold dilutions were spotted onto ONA (~16 h incubation) or SMG (~36 h incubation) agar plates.

### Immunoblotting

Cells were cultured in 150 ml of minimal media (SMG) at 37°C until mid-exponential phase (OD=0.022-0.025). Pellets were collected by filtration, washed and resuspended in 1 ml PBSG (PBS supplemented with 0.1% glycerol). 1.25 OD units of each sample was resuspended in 50 μl PBS containing 400 units of ReadyLyse lysosome (Epicentre), 12.5 units benzonase (Sigma) and a protease-inhibitor cocktail (PIC, Sigma) and incubated for 30 minutes at 37°C. Next, 4x Loading Dye containing DTT (200 mM final) was added and samples were incubated for 5 minutes at 95°C. Protein extracts from tested strains were mixed with Δ*scpB* or Δ*smc* extracts as follows: tested strain only, 1:1 volume of tested strain with Δ, 1:4, 1:6. 5ul of mixed protein extracts were run on Novex™ WedgeWell™ 4 to 12%, Tris-Glycine gels in 1x Laemmli buffer.

Proteins were transferred onto a PVDF membrane (Immobilon-P, Merck Millipore) using wet transfer. Membranes were blocked with 5% (w/v) milk powder in TBS with 0.05% Tween20. 1:2000 or 1:5000 dilutions of rabbit polyclonal sera against *B. subtilis* ScpB or Smc were used as primary antibodies for immunoblotting, respectively. The membrane was developed with HRP-coupled secondary antibodies and chemiluminescence (Amersham ECL Western Blotting Detection Reagent) and visualized on a FUSION FX7 (Vilber).

### Chromatin Immunoprecipitation (ChIP)

ChIP samples were prepared as described previously (Bürmann et al., 2017). Briefly, cells were cultured in 200 ml of minimal media (SMG) at 37°C until mid-exponential phase (OD=0.022-0.030) and fixed with buffer F (50 mM Tris-HCl pH 7.4, 100 mM NaCl, 0.5 mM EGTA pH 8.0, 1 mM EDTA pH 8.0, 10% (w/v) formaldehyde) for 30 minutes at RT with occasional shaking. Cells were harvested by filtration and washed in PBS. Each sample was adjusted for 2 OD units (2 ml at OD600 = 1) and resuspended in TSEMS lysis buffer (50 mM Tris pH 7.4, 50 mM NaCl, 10 mM EDTA pH 8.0, 0.5 M sucrose and PIC (Sigma), 6 mg/ml lysozyme from chicken egg white (Sigma)). After 30 minutes of incubation at 37°C with vigorous shaking protoplasts were washed again in 2 ml TSEMS, resuspended in 1 ml TSEMS, split into 3 aliquots, pelleted and, after flash freezing, stored at −80°C until further use. For time-course experiments 1 l preculture was first grown until mid-exponential phase (OD = 0.022-0.030) and next, appropriate culture volumes were added to fresh pre-warmed SMG so that at given time points 200 ml of culture at mid-exponential could be processed. The cultures were induced with 2 mM theophylline (*Ptheo* promoter). Due to characteristics of the theophylline switch, the pre-culture as well as induction was performed at 30°C.

For ChIP-qPCR, each pellet was resuspended in 2 ml of buffer L (50 mM HEPES-KOH pH 7.5, 140 mM NaCl, 1 mM EDTA pH 8.0, 1% (v/v) Triton X-100, 0.1% (w/v) Na-deoxycholate, 0.1 mg/ml RNaseA and PIC (Sigma)) and transferred to 5 ml round-bottom tubes. Cell suspensions were sonicated 3 x 20 sec on a Bandelin Sonoplus with a MS72 tip (90% pulse and 35% power output). Next, lysates were transferred into 2 ml tubes and centrifuged 10 minutes at 21000g at 4°C. 800 μl of supernatant was used for IP and 200 μl was kept as WCE (whole cell extract).

For IP, first, antibody serum was incubated with Protein G coupled Dynabeads (Invitrogen) in 1:1 ratio for 2.5 hours at 4°C with rotation. Next, beads were washed in buffer L and 50 μl were aliquoted to each sample tube. Samples were incubated for 2 hours at 4°C with rotation, followed by a series of washes with buffer L, buffer L5 (buffer L containing 500 mM NaCl), buffer W (10 mM Tris-HCl pH 8.0, 250 mM LiCl, 0.5% (v/v) NP-40, 0.5% (w/v) sodium deoxycholate, 1 mM EDTA pH 8.0) and buffer TE (10 mM Tris-HCl pH 8.0, 1 mM EDTA pH 8.0). Finally, the beads were resuspended in 520 μl buffer TES (50 mM Tris-HCl pH 8.0, 10 mM EDTA pH 8.0, 1% (w/v) SDS). 300 μl of TES and 20 μl of 10% SDS were also added to WCE. Both tubes were incubated O/N at 65°C with vigorous shaking to reverse formaldehyde crosslinks.

Phenol-chloroform extraction was performed to purify the decrosslinked DNA. Samples were transferred to screw cap 1.5-ml tubes and first mixed vigorously with 500 μl of phenol equilibrated with buffer (10 mM Tris-HCl pH 8.0, 1 mM EDTA). After centrifugation (10 minutes, RT, 13000 rpm), 450 μl of the aqueous phase was transferred to a new screw cap tube and mixed with equal volume of chloroform, followed by centrifugation. 400ul of aqueous phase was recovered for DNA precipitation with 2.5x volume of 100 % ethanol, 0.1x volume of 3M NaOAc and 1.2 μl of GlycoBlue and incubated for 20 minutes at −20°C. Next, samples were centrifuged for 10 minutes at 20000 g at RT and pellets obtained pellets were resuspended in 10 μl of EB (Qiagen) shaking at 55°C for 10 minutes and finally purified with a PCR purification kit, eluting in 50 uL EB.

For qPCR, 1:10 and 1:1000 dilutions in water of IP and WCE were prepared, respectively. Each 10 μl reaction was prepared in duplicate (5 μl Takyon SYBR MasterMix, 1 μl 3 uM primer pair, 4 μl of DNA) and run in Rotor-Gene Q machine (Qiagen). Primer sequences are listed in the Table 2. Data was analysed using PCR Miner server (http://ewindup.info) (Zhao & Fernald, 2005).

**Table 2.**
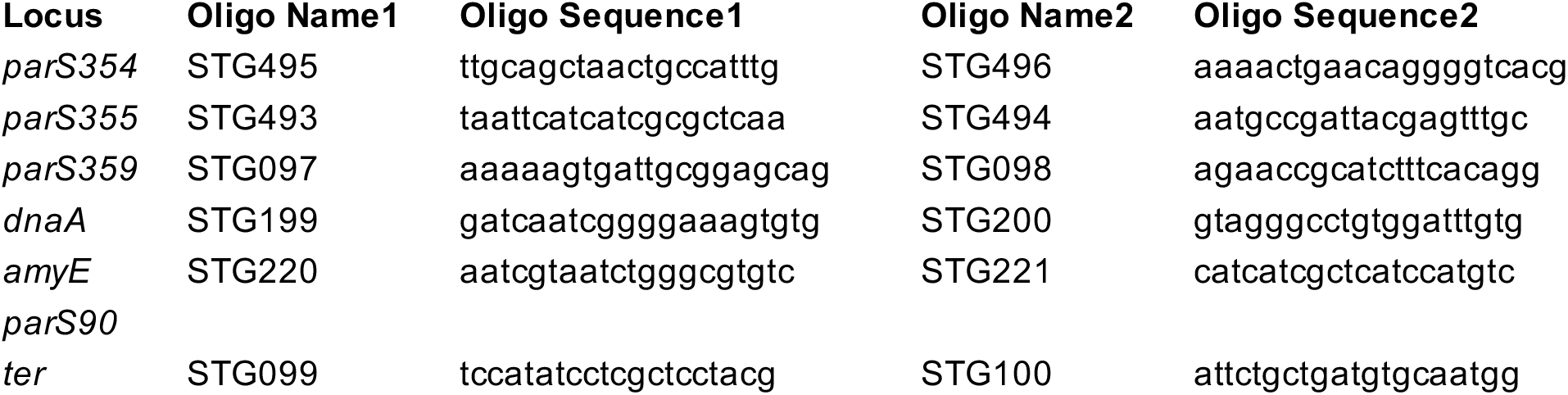
List of oligos used for qPCR

For IP of samples for ChIP-seq, the procedure was the same as for ChIP-qPCR, except for resuspending the pellets in 1 ml of buffer L and sonication in a Covaris E220 water bath sonicator for 5 minutes at 4°C, 100 W, 200 cycles, 10% load and water level 5.

For deep sequencing, the DNA libraries were prepared by Genomic Facility at CIG, UNIL, Lausanne. Briefly, the DNA was fragmented by sonication (Covaris S2) to fragment sizes ranging from 220–250 bp. DNA libraries were prepared using the Ovation Ultralow Library Systems V2 Kit (NuGEN) including 15 cycles of PCR amplification. Five to ten million single end sequence reads were obtained on a HiSeq2500 (Illumina) with 150-bp read length.

### Processing of ChIP-seq reads

Reads were mapped to *Bacillus subtilis* genome NC_000964.3 (for 1A700) or NC_0022898 (for PY79) with bowtie2 using -very-sensitive-local mode. Subsequent data analysis was performed using Seqmonk (http://www.bioinformatics.babraham.ac.uk/projects/seqmonk/) and R. The bin size used is 1 kb. For the enrichment plots the data was smoothened using Local Polynomial Regression Fitting (loess).

### Generation of chromosome conformation capture (3C) libraries

3C libraries were prepared as previously described (Marbouty et al., 2015). Minimal media (SMG) was used instead of LB. Briefly, cells were grown in 400 ml of SMG medium to exponential phase (OD = 0.022-0.030) and fixed with fresh formaldehyde (3% final concentration) for 30 minutes at RT, followed by 30 minutes at 4°C and quenched for 30 minutes with 0.25 M glycine at 4°C. Fixed cells were harvested by filtering, washed in fresh SMG, frozen in liquid nitrogen and stored at −80°C until further use.

For time-course experiments 2 l preculture was first grown until mid-exponential phase (OD = 0.022-0.030) and next, appropriate culture volumes were added to fresh pre-warmed SMG so that at given time points 2 x 200 ml of culture at mid-exponential could be collected (two technical replicates). The cultures were induced with 2 mM theophylline or 1 mM IPTG, depending on the promoter used, *Ptheo* or *Pspank*, respectively. Due to characteristics of the theophylline switch, the pre-culture as well as induction was performed at 30°C.

Frozen pellets were resuspended in 600 μl 1x TE and incubated at RT for 20 minutes with 4 μl of Ready-lyze lysozyme (35U/ul, Tebu Bio). Next, SDS was added to a final concentration of 0.5% and incubated at RT for 10 minutes.

50 μl of lysed cells were aliquoted to 8 tubes containing 450 μl of digestion mix (1x NEB 1 buffer, 1% triton X-100, and 100 U HpaII enzyme (NEB)) and incubated at 37°C for 3 hours with constant shaking. Digested DNA was collected by centrifugation, diluted into 4 tubes containing 8 ml of ligation mix (1x ligation buffer: 50 mM Tris-HCl, 10 mM MgCl2, 10 mM DTT), 1 mM ATP, 0.1 mg/ml BSA, 125 U T4 DNA ligase 5 U/ml) and incubated at 16°C for 4 hours. Ligation reaction was followed by O/N decrosslinking at 65°C in the presence of 250 μg/ml proteinase K (Eurobio) and 5 mM EDTA.

DNA was precipitated with 1 volume of isopropanol and 0.1 Volume of 3 M sodium acetate (pH 5.2, Sigma) at −80°C for 1 hour. After centrifugation, the DNA pellet was resuspended in 1x TE at 30°C for 20 minutes. Next, DNA was extracted once with 400 μl phenol-chloroform-isoamyl alcohol solution and precipitated with 1.5 volume cold 100% ethanol in the presence of 0.1 volume 3 M sodium acetate at −80°C for 30 minutes. The pellet was collected and resuspended in 30 μl TE with RNaseA at 37°C for 30 minutes. All tubes were pooled and the resulting 3C library was quantified on gel using ImageJ.

### Processing of libraries for Illumina sequencing

1 ug of 3C library was suspended in water (final volume 130 μl) and sonicated using Covaris S220 (following manufacturers recommendations to obtain 500 bp target size). Next, DNA was purified with Qiagen PCR purification kit, eluted in 40 μl EB and quantified using NanoDrop. 1 ug of DNA was processed according to manufacturer instructions (Paired-End DNA sample Prep Kit – Illumina – PE-930-1001), except that DNA was ligated to custom-made adapters for 4 hours at RT, followed by inactivation step at 65°C for 20 minutes. DNA was purified with 0.75x AMPure beads and 3 μl were used for 50 μl PCR reaction (12 cycles). Amplified libraries were purified on Qiagen columns and pair end sequenced on an Illumina platform (HiSeq4000 or NextSeq).

### Processing of PE reads and generation of contact maps

Sequencing data was demultiplexed, adapters trimmed, and PCR duplicates removed using custom scripts. Next, data was processed as described at https://github.com/axelcournac/3C_tutorial. Briefly, bowtie2 in -very sensitive-local mode was used for mapping for each mate. After sorting and merging both mates, the reads of mapping quality >30 were filtered out and assigned to a restriction fragment. Uninformative events like recircularization on itself (loops), uncut fragments, re-ligations in original orientation were discarded (Cournac et al., 2012) and only pairs of reads corresponding to long-range interactions were used for generation of contact maps (between 5-8% of all reads). The bin size used is 10 kb. Next, contact maps were normalized through the sequential component normalization procedure (SCN, (Cournac et al., 2012)). Subsequent visualization was done using MATLAB (R2019b). To facilitate visualization of the contact matrices, first we applied to the SCN matrices the log10 and then a Gaussian filter (H=1) to smooth the image. The scale bar next to the maps represent the contact frequencies in log_10_– the darker the colour, the higher the frequency of contacts between given loci.

## Acknowledgements

We thank Hugo Brandao, Leonid Mirny and Xindan Wang for sharing unpublished data and comments on the manuscript. We are grateful to Frank Bürmann and all members of the Gruber laboratory for stimulating discussions and critical feedback. We thank Marc Garcia-Garcera and Bjorn Vessman for continuous support with R and Python and the Genomic Technologies Facility (GTF, UNIL, Lausanne) and Next Generation Sequencing Core Facility (CNRS, I2BC, Gif-sur-Yvette) for ChIP-seq library preparation and deep sequencing.

---

**Figure S1.**
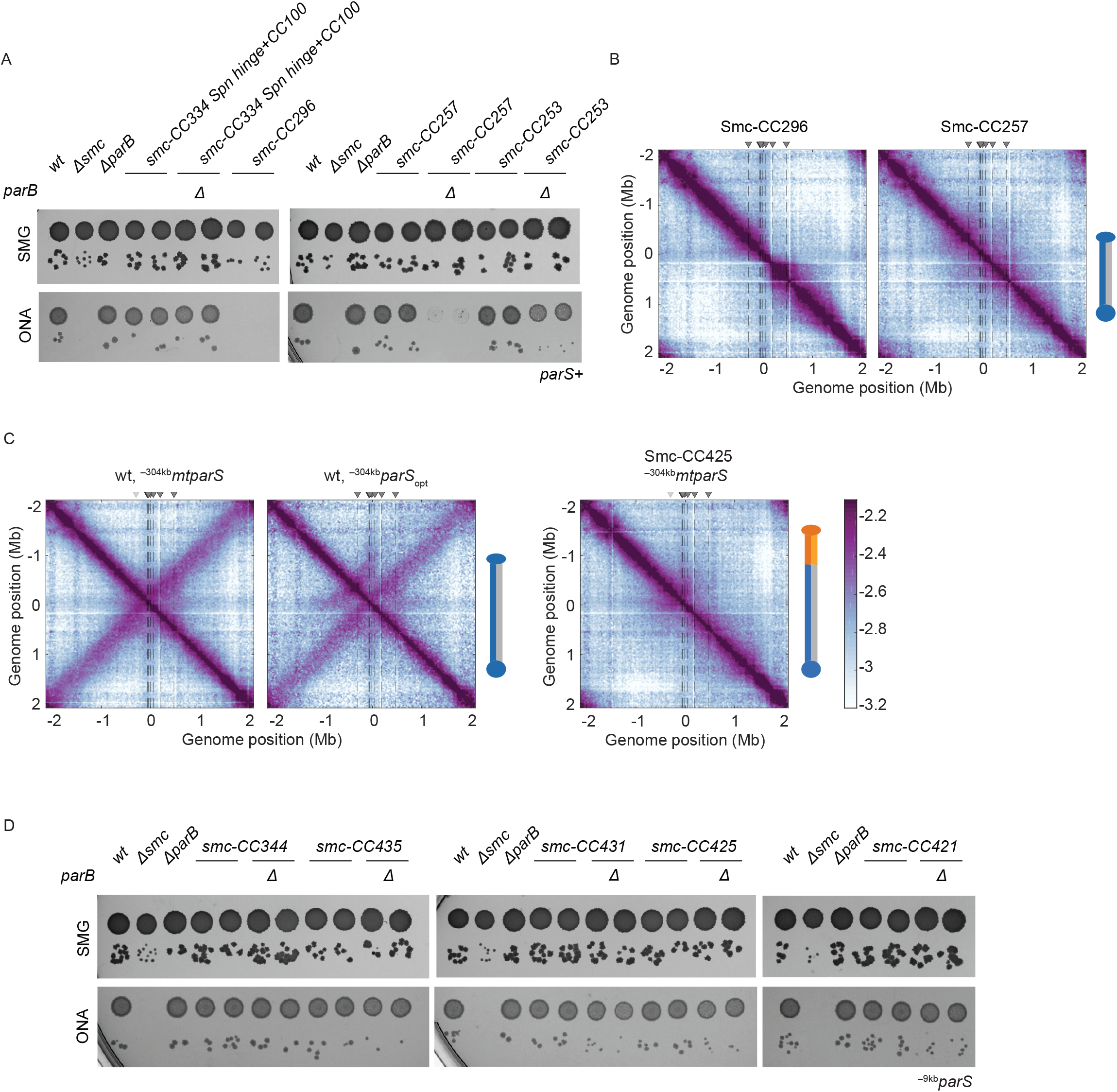
Different variants of Smc proteins and their ability to support growth and chromosome alignment. A. Spotting assay of strains with modified Smc coiled coil in wild type or sensitised background (Δ*parB*). Prepared as described in Figure 1B. Two clones for each tested mutant were spotted. Combining Δ*parB* with Smc-CC296 was unsuccessful. *Spn* hinge + CC100, *Streptococcus pneumoniae* hinge and 100 amino acids of hinge-proximal coiled coil. B. Normalized 3C-seq contact maps for strains carrying all the wild type *parS* sites and Smc’s with shortened coiled coils that were dead (*left panel*) or sick on rich medium (*right panel*). C. Normalized 3C-seq contact maps for strains with inactivated *parS-334* or *parS-334* mutated to *parS-359* sequence (*parS*_*opt*_) and wild type Smc (*left and middle panel*) or Smc-CC425 (*right panel*). D. Spotting assay of strains with elongated Smc coiled coil in single ^−9k*b*^*parS*_*opt*_ (*parS-359*) background with or without *parB*. Prepared as described in Figure 1B. Two clones for each tested mutant were spotted.

**Figure S2.**
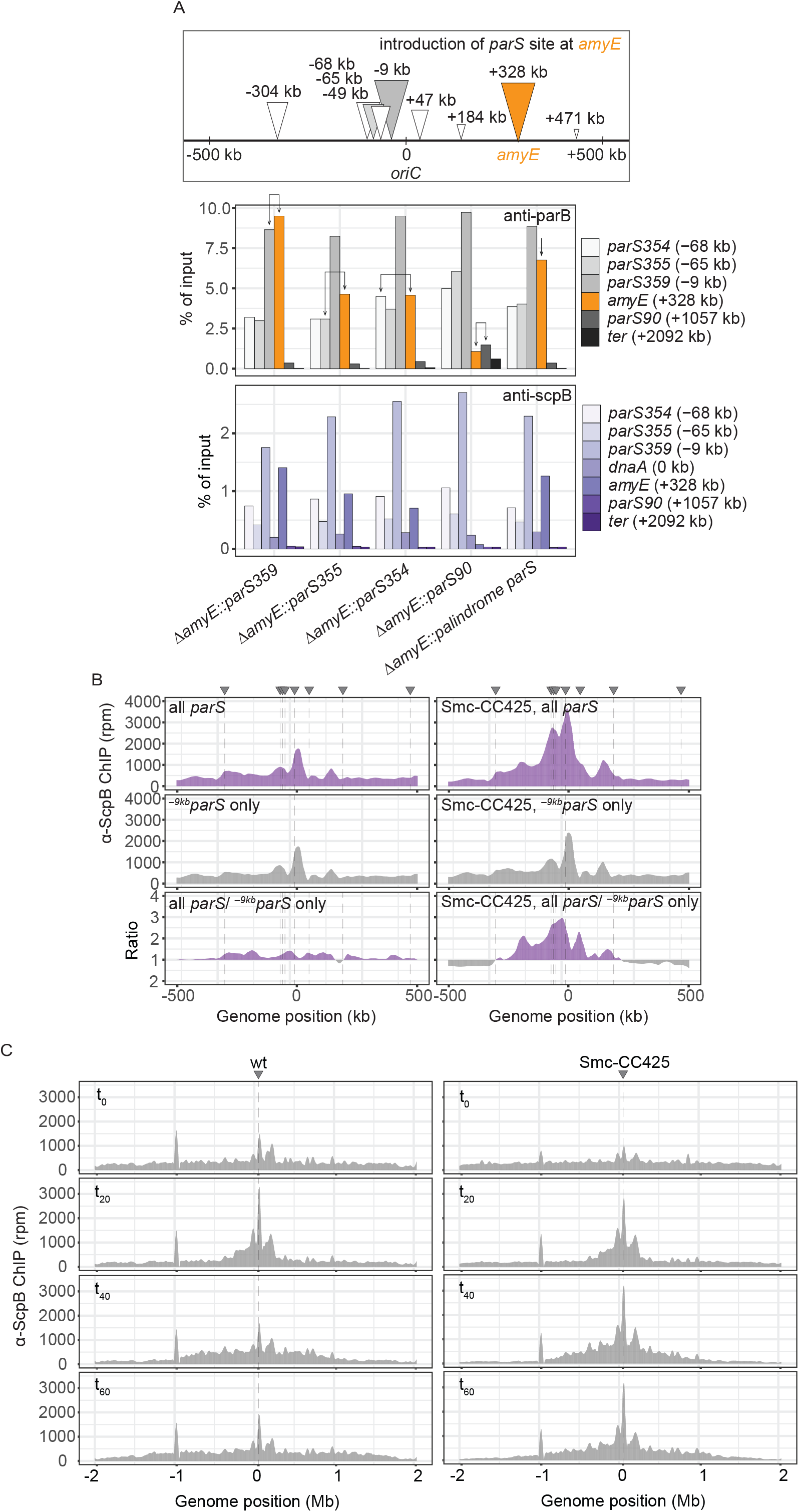
^−9k*b*^*parS* (*parS-359*) sequence recruits ParB most efficiently regardless the position. A. Scheme depicting the experimental set up: different wild type *parS* sites as well as a perfect palindromic *parS* site were engineered at *amyE* locus (orange) in an otherwise wild type background (*top panel*). Chromatin-immunoprecipitation coupled to quantitative PCR (ChIP-qPCR) using α-ParB and α-ScpB serum to test for recruitment of ParB (*middle panel*) and subsequent recruitment of Smc (*bottom panel*). Recruitment of ParB to endogenous *parS* and engineered *parS* sites (orange) are indicated with arrows for each strain. The enrichment observed by ectopic integration at *amyE* were similar to one seen at the respective endogenous loci demonstrating that the *parS* sequence determines the efficiency of ParB recruitment largely irrespective of the genomic neighborhood. The results also supported the notion that the strength of natural *parS* sequences generally decreased with distance from the replication origin as also indicated by ChIP-seq (Figure 1A) (Minnen et al., 2016) and ChIP-chip (Breier & Grossman, 2007). B. Close up of ChIP-seq experiment shown in Figure 1A and 1B. Lines indicate *parS* positions. C. Read count distribution for chromatin immunoprecipitation coupled to deep sequencing (ChIP-seq) using α-ScpB serum for the time course presented in Figure 2C. *Left panel*, strain carrying wild-type Smc. *Right panel*, strain carrying elongated Smc (Smc-CC425). Strains harbour a single loading site, ^−*9kb*^*parS*_opt_ (*parS-359*), and theophylline-inducible *parB* gene. Triangles indicate positions of parS sites: light grey triangle is weak parS at +1058 kb, orange triangle is *parS*_opt_ site introduced at *amyE* locus. Dashed lines correspond to active *parS* site(s) in given experiment.

**Figure S3.**
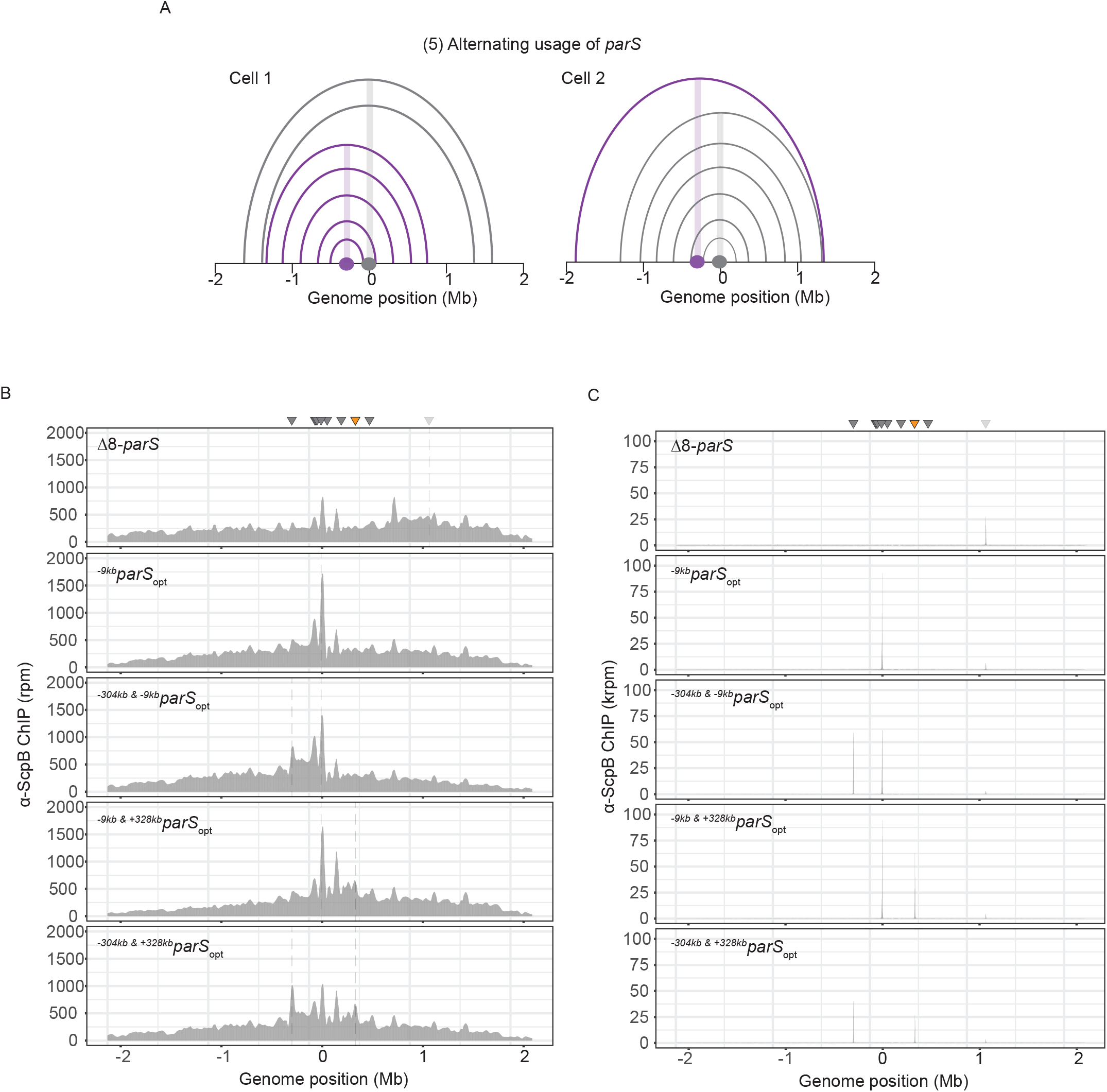
Wild-type Smc protein generates overlapping chromosome folding patterns A. Schemes depicting possible scenario for collision avoidance and collision resolution: exclusive usage of a single *parS* site by SMC complexes as a consequence of temporary inactivation of remaining *parS* sites. B. Read count distribution for chromatin immunoprecipitation coupled to deep sequencing (ChIP-seq) using α-ScpB serum for a strain with eight *parS* sites deleted, single *parS*_opt_ at −9kb, two *parS* sites at −304kb and −9kb, −9kb and +328kb or −304kb and +328kb. Represented as in Figure S2C. C. Read count distribution for chromatin immunoprecipitation coupled to deep sequencing (ChIP-seq) using α-ParB serum for respective strains in S3B.

**Figure S4.**
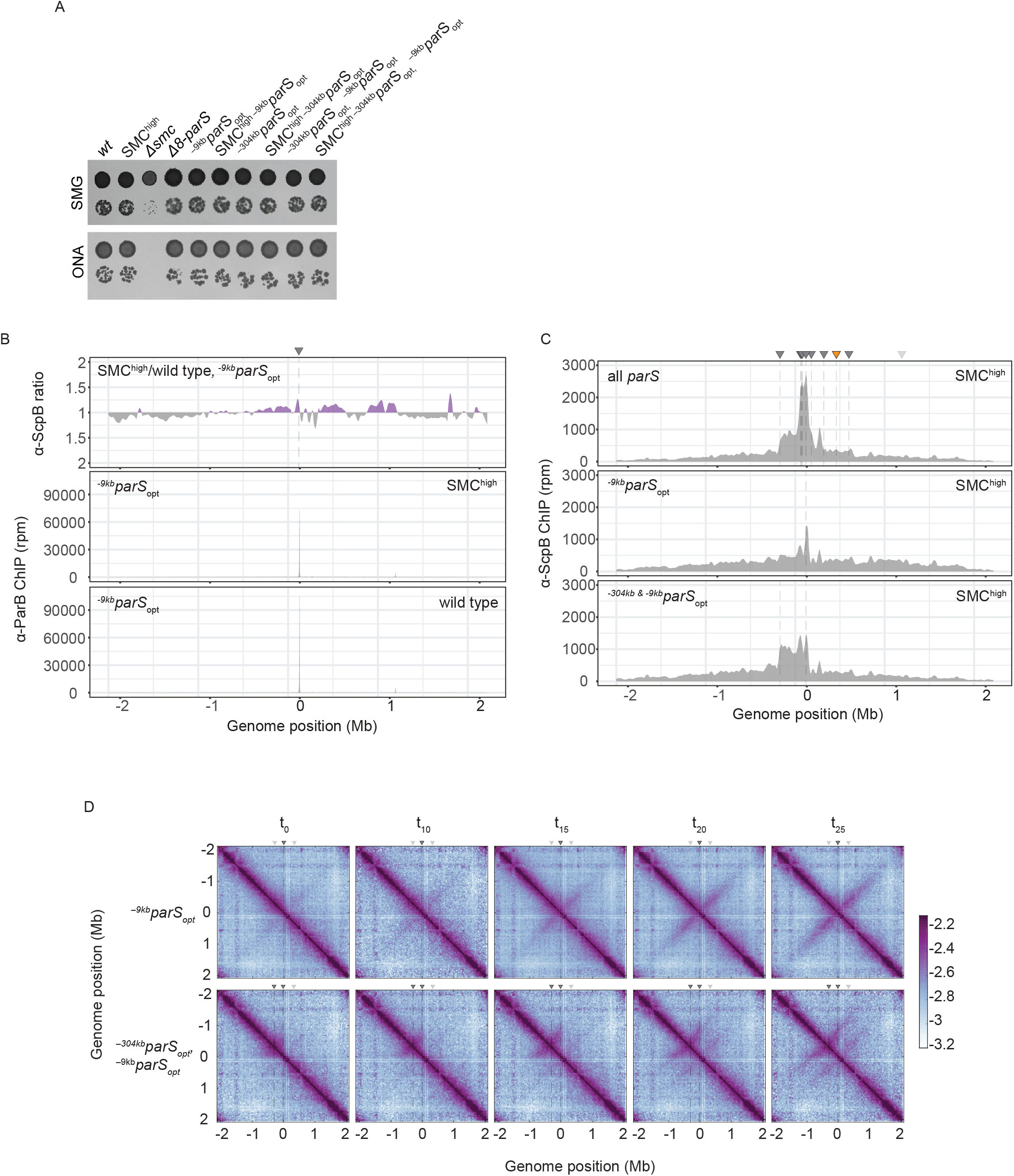
Synchronized loading of SMC hampers chromosome organization. A. Spotting assay comparing strains with increased amount of Smc vs wild type with respective number of *parS*_*opt*_ sites. Prepared as described in Figure 1B B. Ratio plots for ChIP-seq read counts comparing SMC^high^ strain and wild-type levels of Smc with a single *parS* site (^−9*kb*^parS_opt_). Representation as in Figure 2B (*top panel*). Read count for α-ParB ChIP-seq in the same strains (*middle and bottom panels*). C. α-ScpB ChIP-seq read counts for SMC^high^ strains with all *parS* sites present, single *parS*_opt_ at −9kb and two *parS* sites at −304kb and −9kb. Represented as in Figure S3B. D. Normalized 3C-seq contact maps for time course experiment for strains with a single loading site *parS*_*opt*_ (at −9kb, *top row*) or two *parS*_*opt*_ (at −304kb and −9kb, *bottom row*) and IPTG inducible ParB.

## References

Banigan, E. J., van den Berg, A. A., Brandão, H. B., Marko, J. F., & Mirny, L. A. (2020). Chromosome organization by one-sided and two-sided loop extrusion. ELife, 9, 1–46. https://doi.org/10.7554/eLife.53558

Böhm, K., Giacomelli, G., Schmidt, A., Imhof, A., Koszul, R., Marbouty, M., & Bramkamp, M. (2020). Chromosome organization by a conserved condensin-ParB system in the actinobacterium Corynebacterium glutamicum. Nature Communications, 11(1), 1485. https://doi.org/10.1038/s41467-020-15238-4

Brandão, H. B., Paul, P., van den Berg, A. A., Rudner, D. Z., Wang, X., & Mirny, L. A. (2019). RNA polymerases as moving barriers to condensin loop extrusion. Proceedings of the National Academy of Sciences of the United States of America, 116(41), 20489–20499. https://doi.org/10.1073/pnas.1907009116

Brandão, H. B., Ren, Z., Karaboja, X., Mirny, L. A., & Wang, X. (2020). DNA-loop extruding SMC complexes can traverse one another *in vivo* BioRxiv, 2020.10.26.356329. https://doi.org/10.1101/2020.10.26.356329

Brandão, H. B., Wang, X., Paul, P., Berg, A. A. Van Den, David, Z., & Mirny, L. A. (2019). RNA polymerases as moving barriers to condensin loop extrusion Abstract : 1.

Breier, A. M., & Grossman, A. D. (2007). Whole-genome analysis of the chromosome partitioning and sporulation protein Spo0J (ParB) reveals spreading and origin-distal sites on the Bacillus subtilis chromosome. Molecular Microbiology, 64(3), 703–718. https://doi.org/10.1111/j.1365-2958.2007.05690.x

Bürmann, F., Basfeld, A., Vazquez Nunez, R., Diebold-Durand, M. L., Wilhelm, L., & Gruber, S. (2017). Tuned SMC Arms Drive Chromosomal Loading of Prokaryotic Condensin. Molecular Cell, 65(5), 861–872.e9. https://doi.org/10.1016/j.molcel.2017.01.026

Bürmann, F., & Gruber, S. (2015). SMC condensin: promoting cohesion of replicon arms. Nature Structural & Molecular Biology, 22(9), 653–655. https://doi.org/10.1038/nsmb.3082

Bürmann, F., Shin, H. C., Basquin, J., Soh, Y. M., Giménez-Oya, V., Kim, Y. G., Oh, B. H., & Gruber, S. (2013). An asymmetric SMC-kleisin bridge in prokaryotic condensin. Nature Structural and Molecular Biology, 20(3), 371–379. https://doi.org/10.1038/nsmb.2488

Cournac, A., Marie-Nelly, H., Marbouty, M., Koszul, R., & Mozziconacci, J. (2012). Normalization of a chromosomal contact map. BMC Genomics, 13(1). https://doi.org/10.1186/1471-2164-13-436

Davidson, I. F., Bauer, B., Goetz, D., Tang, W., Wutz, G., & Peters, J.-M. (2019). DNA loop extrusion by human cohesin. Science, 366(6471), 1338 LP – 1345. https://doi.org/10.1126/science.aaz3418

Dyson, S., Segura, J., Martínez-garcía, B., Valdés, A., & Roca, J. (2020). Condensin minimizes topoisomerase II-mediated entanglements of DNA in vivo. 1–14. https://doi.org/10.15252/embj.2020105393

Ganji, M., Shaltiel, I. A., Bisht, S., Kim, E., Kalichava, A., Haering, C. H., & Dekker, C. (2018). Real-time imaging of DNA loop extrusion by condensin. Science, 360(6384), 102–105. https://doi.org/10.1126/science.aar7831

Gligoris, T. G., Scheinost, J. C., Bürmann, F., Petela, N., Chan, K. L., Uluocak, P., Beckouët, F., Gruber, S., Nasmyth, K., & Löwe, J. (2014). Closing the cohesin ring: Structure and function of its Smc3-kleisin interface. Science, 346(6212), 963–967. https://doi.org/10.1126/science.1256917

Graham, T. G. W., Wang, X., Song, D., Etson, C. M., van Oijen, A. M., Rudner, D. Z., & Loparo, J. J. (2014). ParB spreading requires DNA bridging. Genes and Development, 28(11), 1228–1238. https://doi.org/10.1101/gad.242206.114

Gruber, S., Arumugam, P., Katou, Y., Kuglitsch, D., Helmhart, W., Shirahige, K., & Nasmyth, K. (2006). Evidence that Loading of Cohesin Onto Chromosomes Involves Opening of Its SMC Hinge. Cell, 127(3), 523–537. https://doi.org/10.1016/j.cell.2006.08.048

Gruber, S., & Errington, J. (2009). Recruitment of Condensin to Replication Origin Regions by ParB/SpoOJ Promotes Chromosome Segregation in B. subtilis. Cell, 137(4), 685–696. https://doi.org/10.1016/j.cell.2009.02.035

Gruber, S., Veening, J. W., Bach, J., Blettinger, M., Bramkamp, M., & Errington, J. (2014). Interlinked sister chromosomes arise in the absence of condensin during fast replication in B. subtilis. Current Biology, 24(3), 293–298. https://doi.org/10.1016/j.cub.2013.12.049

Jalal, A. S. B., Tran, N. T., & Le, T. B. K. (2020). ParB spreading on DNA requires cytidine triphosphate in vitro. ELife, 9, e53515. https://doi.org/10.7554/eLife.53515

Kim, E., Kerssemakers, J., Shaltiel, I. A., Haering, C. H., & Dekker, C. (2020). DNA-loop extruding condensin complexes can traverse one another. Nature, 579(7799), 438–442. https://doi.org/10.1038/s41586-020-2067-5

Kim, Y., Shi, Z., Zhang, H., Finkelstein, I. J., & Yu, H. (2019). Human cohesin compacts DNA by loop extrusion. Science, 366(6471), 1345–1349. https://doi.org/10.1126/science.aaz4475

Ku, B., Lim, J. H., Shin, H. C., Shin, S. Y., & Oh, B. H. (2010). Crystal structure of the MukB hinge domain with coiled-coil stretches and its functional implications. Proteins: Structure, Function and Bioinformatics, 78(6), 1483–1490. https://doi.org/10.1002/prot.22664

Lagage, V., Boccard, F., & Vallet-Gely, I. (2016). Regional Control of Chromosome Segregation in Pseudomonas aeruginosa. PLoS Genetics, 12(11), e1006428. https://doi.org/10.1371/journal.pgen.1006428

Lee, P. S., Lin, D. C. H., Moriya, S., & Grossman, A. D. (2003). Effects of the chromosome partitioning protein Spo0J (ParB) on oriC positioning and replication initiation in Bacillus subtilis. Journal of Bacteriology, 185(4), 1326–1337. https://doi.org/10.1128/JB.185.4.1326-1337.2003

Lioy, V. S., Cournac, A., Marbouty, M., Duigou, S., Mozziconacci, J., Espéli, O., Boccard, F., & Koszul, R. (2018). Multiscale Structuring of the E. coli Chromosome by Nucleoid-Associated and Condensin Proteins. Cell, 771–783. https://doi.org/10.1016/j.cell.2017.12.027

Lioy, V. S., Junier, I., Lagage, V., Vallet, I., & Boccard, F. (2020). Distinct Activities of Bacterial Condensins for Chromosome Management in Pseudomonas aeruginosa. Cell Reports, 33(5). https://doi.org/10.1016/j.celrep.2020.108344

Livny, J., Yamaichi, Y., & Waldor, M. K. (2007). Distribution of centromere-like parS sites in bacteria: Insights from comparative genomics. Journal of Bacteriology, 189(23), 8693–8703. https://doi.org/10.1128/JB.01239-07

Mäkelä, J., & Sherratt, D. J. (2020). Organization of the Escherichia coli Chromosome by a MukBEF Axial Core. Molecular Cell, 78(2), 250–260.e5. https://doi.org/10.1016/j.molcel.2020.02.003

Marbouty, M., Le Gall, A., Cattoni, D. I., Cournac, A., Koh, A., Fiche, J. B., Mozziconacci, J., Murray, H., Koszul, R., & Nollmann, M. (2015). Condensin- and Replication-Mediated Bacterial Chromosome Folding and Origin Condensation Revealed by Hi-C and Super-resolution Imaging. Molecular Cell, 59(4), 588–602. https://doi.org/10.1016/j.molcel.2015.07.020

Minnen, A., Attaiech, L., Thon, M., Gruber, S., & Veening, J.-W. (2011). SMC is recruited to oriC by ParB and promotes chromosome segregation in Streptococcus pneumoniae. Molecular Microbiology, 81(3), 676–688. https://doi.org/10.1111/j.1365-2958.2011.07722.x

Minnen, A., Bürmann, F., Wilhelm, L., Anchimiuk, A., Diebold-Durand, M. L., & Gruber, S. (2016). Control of Smc Coiled Coil Architecture by the ATPase Heads Facilitates Targeting to Chromosomal ParB/parS and Release onto Flanking DNA. Cell Reports, 14(8), 2003–2016. https://doi.org/10.1016/j.celrep.2016.01.066

Orlandini, E., Marenduzzo, D., & Michieletto, D. (2019). Synergy of topoisomerase and structural-maintenance-of-chromosomes proteins creates a universal pathway to simplify genome topology. Proceedings of the National Academy of Sciences of the United States of America, 116(17), 8149–8154. https://doi.org/10.1073/pnas.1815394116

Osorio-Valeriano, M., Altegoer, F., Steinchen, W., Urban, S., Liu, Y., Bange, G., & Thanbichler, M. (2019). ParB-type DNA Segregation Proteins Are CTP-Dependent Molecular Switches. Cell, 179(7), 1512–1524.e15. https://doi.org/10.1016/j.cell.2019.11.015

Racko, D., Benedetti, F., Goundaroulis, D., & Stasiak, A. (2018). Chromatin loop extrusion and chromatin unknotting. Polymers, 10(10), 1–11. https://doi.org/10.3390/polym10101126

Soh, Y.-M., Davidson, I. F., Zamuner, S., Basquin, J., Bock, F. P., Taschner, M., Veening, J.-W., De Los Rios, P., Peters, J.-M., & Gruber, S. (2019). Self-organization of parS centromeres by the ParB CTP hydrolase. Science, 366(6469), 1129–1133. https://doi.org/10.1126/science.aay3965

Srinivasan, M., Scheinost, J. C., Petela, N. J., Gligoris, T. G., Wissler, M., Ogushi, S., Collier, J. E., Voulgaris, M., Kurze, A., Chan, K. L., Hu, B., Costanzo, V., & Nasmyth, K. A. (2018). The Cohesin Ring Uses Its Hinge to Organize DNA Using Non-topological as well as Topological Mechanisms. Cell, 173(6), 1508–1519.e18. https://doi.org/10.1016/j.cell.2018.04.015

Sullivan, N. L., Marquis, K. A., & Rudner, D. Z. (2009). Recruitment of SMC by ParB-parS Organizes the Origin Region and Promotes Efficient Chromosome Segregation. Cell, 137(4), 697–707. https://doi.org/10.1016/j.cell.2009.04.044

Tran, N. T., Laub, M. T., & Le, T. B. K. (2017). SMC Progressively Aligns Chromosomal Arms in Caulobacter crescentus but Is Antagonized by Convergent Transcription. Cell Reports, 20(9), 2057–2071. https://doi.org/10.1016/j.celrep.2017.08.026

Wang, X., Brandão, H. B., Le, T. B. K., Laub, M. T., & Rudner, D. Z. (2017). Bacillus subtilis SMC complexes juxtapose chromosome arms as they travel from origin to terminus. Science, 355(6324), 1–32. https://doi.org/10.1126/science.aai8982

Wang, X., Le, T. B. K., Lajoie, B. R., Dekker, J., Laub, M. T., & Rudner, D. Z. (2015). Condensin promotes the juxtaposition of dna flanking its loading site in Bacillus subtilis. Genes and Development, 29(15), 1661–1675. https://doi.org/10.1101/gad.265876.115

Wang, X., Tang, O. W., Riley, E. P., & Rudner, D. Z. (2014). The SMC condensin complex is required for origin segregation in Bacillus subtilis. Current Biology, 24(3), 287–292. https://doi.org/10.1016/j.cub.2013.11.050

Wilhelm, L., Bürmann, F., Minnen, A., Shin, H. C., Toseland, C. P., Oh, B. H., & Gruber, S. (2015). SMC condensin entraps chromosomal DNA by an ATP hydrolysis dependent loading mechanism in Bacillus subtilis. ELife, 4(MAY), 1–18. https://doi.org/10.7554/eLife.06659

Yatskevich, S., Rhodes, J., & Nasmyth, K. (2019). Organization of Chromosomal DNA by SMC Complexes. Annual Review of Genetics, 53(1). https://doi.org/10.1146/annurev-genet-112618-043633

Zhao, S., & Fernald, R. D. (2005). Comprehensive algorithm for quantitative real-time polymerase chain reaction. Journal of Computational Biology, 12(8), 1047–1064. https://doi.org/10.1089/cmb.2005.12.1047

